# Glutamylation of Npm2 and Nap1 acidic disordered regions increases DNA charge mimicry to enhance chaperone efficiency

**DOI:** 10.1101/2023.09.18.558337

**Authors:** Benjamin M. Lorton, Christopher Warren, Humaira Ilyas, Prithviraj Nandigrami, Subray Hegde, Sean Cahill, Stephanie M Lehman, Jeffrey Shabanowitz, Donald F. Hunt, Andras Fiser, David Cowburn, David Shechter

**Author notes:** Contributed equally (Benjamin M. Lorton, Christopher Warren, Humaira Ilyas).

## Abstract

Histone chaperones–structurally diverse, non-catalytic proteins enriched with acidic intrinsically disordered regions (IDRs)–protect histones from spurious nucleic acid interactions and guide their deposition into and out of nucleosomes. Despite their conservation and ubiquity, the function of the chaperone acidic IDRs remains unclear. Here, we show that the *Xenopus laevis* Npm2 and Nap1 acidic IDRs are substrates for TTLL4 (Tubulin Tyrosine Ligase Like 4)-catalyzed post-translational glutamate-glutamylation. We demonstrate that, to bind, stabilize, and deposit histones into nucleosomes, chaperone acidic IDRs function as DNA mimetics. Our biochemical, computational, and biophysical studies reveal that glutamylation of these chaperone polyelectrolyte acidic stretches functions to enhance DNA electrostatic mimicry, promoting the binding and stabilization of H2A/H2B heterodimers and facilitating nucleosome assembly. This discovery provides insights into both the previously unclear function of the acidic IDRs and the regulatory role of post-translational modifications in chromatin dynamics.

## Introduction

Nucleosomes–the complex of histone proteins and DNA that compacts and regulates the eukaryotic genome–are assembled in a tightly regulated process. Histone chaperones bind to the basic histone proteins (H2A, H2B, H3, H4 and H1 linker histones), promote proper nucleosome assembly, and reduce non-nucleosomal histone:nucleic acid interactions ^1-3^, while ATP-dependent remodelers position the nucleosomes. To coordinate RNA polymerase passage, some chaperones like FACT engage both histones and assembled nucleosomes ^4-6^. NMR analysis of human and *Drosophila* H2A/H2B heterodimers showed that, while the overall structure is similar to nucleosomal H2A/H2B, the histones in solution are dynamic ^7,8^.

Despite performing common functions, the large family of histone chaperones are structurally diverse. As we previously showed, most contain long intrinsically disordered regions (IDRs) that are enriched with acidic stretches ^9,10^. Structures of histone chaperones bound to their cognate histones have revealed common interaction themes, including an “anchoring and capping” mechanism of H2A/H2B-interacting histone chaperone acidic stretches ^11-13^. Others have shown direct interactions and function of acidic IDRs with histones, including: the SPT16 subunit IDR of FACT interaction with H2A/H2B ^5^; MCM2 acidic IDR interaction with Asf1 and H3/H4 ^14,15^; and on octamers by acidic IDRs of APLF ^16^. Studies of the Nap1 chaperone:histone interactions are challenged both by its oligomeric nature and the likely multiple binding modes, but it has been demonstrated that histone interactions are enhanced by the presence of the Nap1 N- and C-terminal acidic IDRs ^17-21^. A recent study further suggests that Nap1 acidic IDRs exhibit a “penetrating, fuzzy binding mechanism” to histones mediated by electrostatic interactions ^22^.

γ-Glutamate-glutamylation is a reversible post-translational modification (PTM) catalyzed by an ATP-dependent addition of a glutamate amino acid to an existing peptidyl glutamate ^23^. The initiator glutamate is added to the γ-carboxyl group of a glutamate sidechain through an isopeptide bond. Additional glutamates can be added either to the α-carboxyl or γ-carboxyl groups of the branch point glutamate, resulting in polyglutamylation. The family of Tubulin Tyrosine Like Ligase (TTLL) enzymes catalyze glutamylation ^23^, while deglutamylation is catalyzed by members of the Cytosolic Carboxypeptidase (CCP) enzyme family ^24^. Glutamylation was first identified on alpha and beta tubulin ^25-27^, where it modulates interactions with microtubule associated proteins, thereby influencing the dynamics of microtubule arrays in cells ^28^. Glutamylation was also identified on histone chaperones, including Nucleophosmin (NPM1) ^29^, Nucleoplasmin (Npm2) ^30^, Nucleosome Assembly Proteins 1 and 2 (Nap1, Nap2), Acidic Nuclear Phosphoprotein 32 E (ANP32E) ^31-33^, and PELP1 ^34^. While one study demonstrated that glutamylation of the C-terminal acidic domain of Nap1 increased deposition of linker histones and was required for condensation of mitotic chromosomes ^31^, how glutamylation influences chaperone function remains largely unknown.

The fundamental mechanisms underlying the transfer of histones from chaperone to DNA remain largely unknown ^35-38^. Recent studies have implicated long-timescale dynamic polyelectrolyte competition in chaperone IDR mediation of H1 linker histone removal ^39^. In light of this and related observations on the cellular function of diffuse and multivalent charge-charge interactions ^40,41^, we suggest that intermediate, electrostatically mediated states are likely critical for the ordered formation and disassembly of nucleosomes. To understand histone chaperone mechanisms, we hypothesized that chaperone acidic IDRs– made more acidic by glutamate-glutamylation–mimic DNA to promote histone stability and enhance histone deposition and removal. In this study, we provide robust computational, biochemical, and biophysical evidence that post-translationally glutamylated histone chaperone acidic IDRs behave as polyelectrolytes to bind to histones. This binding stabilizes H2A/H2B for either nucleosomal deposition or for aggregate removal.

## Results

### Nap1 and Npm2 are glutamate-glutamylated

Due to their abundance in *Xenopus laevis* eggs and their well described and widely conserved function in histone storage and deposition, we focused on the canonical H2A/H2B chaperones Npm2 (also known as Nucleoplasmin) and Nap1 (NAP1L1, Nucleosome Assembly Protein 1 Like 1, here referred to as Nap1). We previously showed that Npm2 from *Xenopus laevis* eggs and oocytes was glutamylated within the A2 acidic disordered stretch^30^ (**Figure 1a,b**). Here, we purified Nap1 from *Xenopus* egg extract and showed that it is post-translationally glutamylated (**Figures 1a,b and Supplementary Figure S1a-e**). As observed by immunoblot and mass spectrometry (**Supplementary Figure S1f**), *Xenopus* Nap1 has two main genetic isoforms (Nap1.S and Nap1.L) and two splice isoforms (X1 and X2). The X1 isoform contains ten additional residues at its C-terminus and harbors a single farnesyl-cysteine that is not present on isoform X2 (**Figure 1c**, highlighted green). In this study, we used the most abundant isoform (X2; **Supplementary Figure S1g**). We identified Nap1 mono-glutamylation within both its N-terminal A1 and C-terminal A3 acidic disordered stretches (**Figure 1c**, highlighted orange). Glutamylation of the Nap1 A1 acidic stretch occurs in distribution (one to six sites, with three most abundant at 30-50% of total protein) (**Figures 1d,f**). The Nap1 A3 acidic patch contained one to eleven sites of glutamylation, with eight sites being the most abundant form at 30-50% of total protein (**Figures 1e,f**). While outside of our Npm2 and Nap1 experiments in this study, we also observed by immunoblot that the egg histone chaperone NASP (formerly known as N1) was glutamylated (**Supplementary Figure S1h,i**).

**Figure 1.**
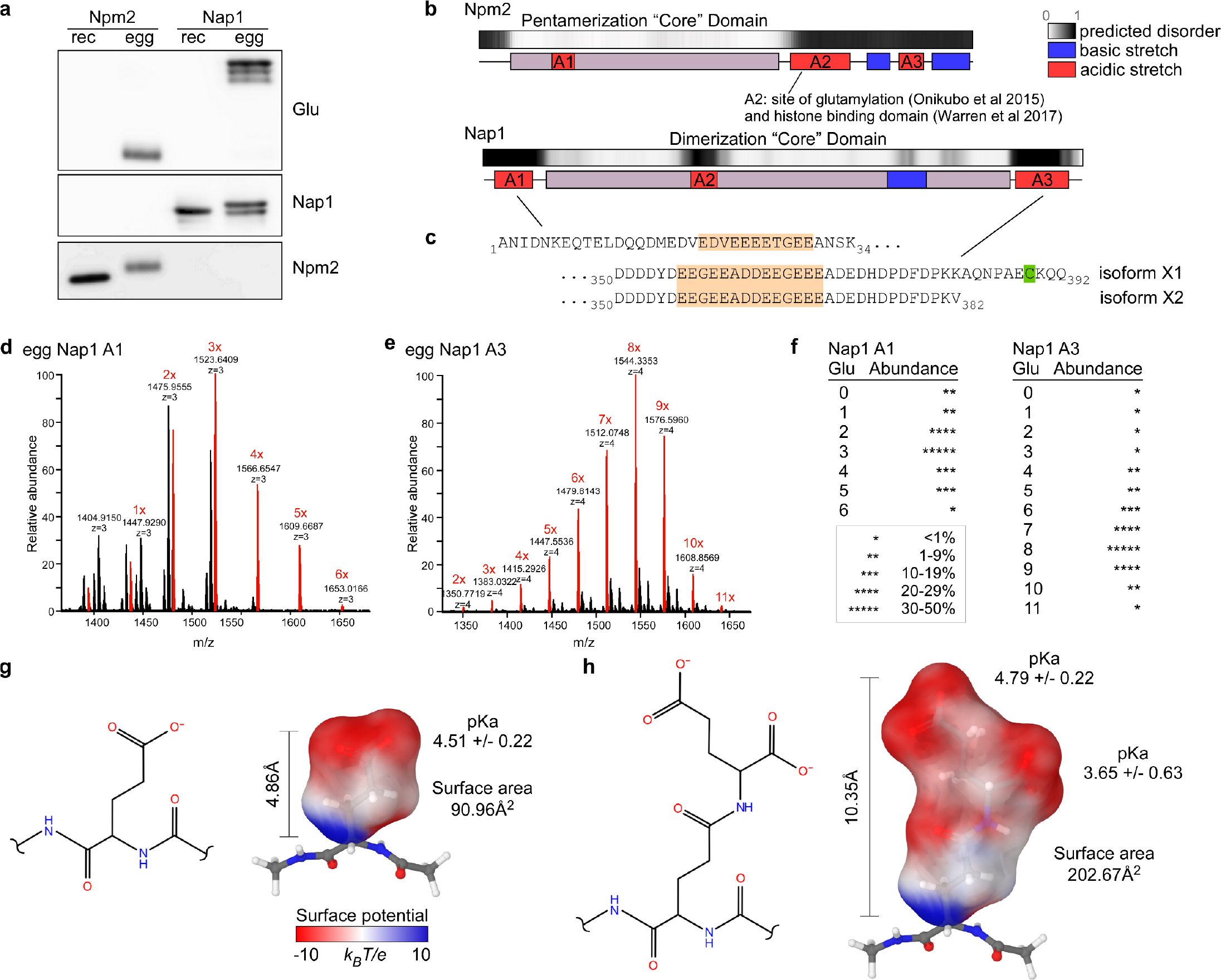
*Xenopus laevis* histone chaperones Npm2 and Nap1 are glutamate-glutamylated on their acidic IDRs. **a**. Recombinant (rec) and purified egg Npm2 and Nap1 were immunoblotted for glutamylation (glu, top), Nap1 (middle), and Npm2 (bottom) showing glutamylation specifically on endogenous chaperones. **b**. Predicted disorder (black and white) and domain maps of Npm2 (top) and Nap1 (bottom) showing acidic (red) and basic (blue) patches coincident with intrinsically disordered tails. **c**. Sites of glutamylation (highlighted orange) detected by mass spectrometry on endogenous Nap1 purified from *Xenopus* eggs. A single farnesyl cysteine (highlighted green) was detected in the longer Nap1 X1 isoform. **d**. Quantification of glutamylation on Nap1 acidic patch A1 detected by mass spectrometry. **e**. Quantification of glutamylation on Nap1 acidic patch A3 detected by mass spectrometry. **f**. Relative abundance of Nap1 A1 and A3 glutamylation determined from mass spectrometry. **g**. Model of a peptidyl glutamate showing stick and electrostatic surface potential representations **h**. Same as (a) but model of post-translational glutamate-glutamylation.

To define the physical nature of monoglutamylation, we computationally modeled both glutamate and γ-glutamylglutamate. The unmodified glutamate sidechain measures 4.86Å in length with a surface area of 91Å^2^, and a predicted p*K*_a_ of 4.51 ± 0.22 for the γ-carboxyl group (**Figure 1g**). The γ-glutamyl-glutamate side chain was more than doubled in length and surface area, measuring 10.35Å and 202.67Å^2^, respectively, and contained an additional negative charge; the carboxyl groups predicted p*K*_a_’s are 4.79 ± 0.22 and 3.65 ± 0.63 (**Figure 1h**). Thus, glutamylation dramatically increased the size and acidic character of the glutamate side chain (**Supplemental Data 1** for models).

### Histone chaperone intrinsically disordered acidic tails are important for histone binding

To test the contributions of the Nap1 intrinsically disordered acidic tails to histone binding, we performed a competitive pulldown assay using H2A.S2/H2B heterodimers (H2A contained a C-terminal StrepII-tag, denoted S2) and Nap1 full-length or truncations lacking either or both A1 or A3 acidic tails (**Figure 2a**). In the pulldowns, all four proteins bound H2A.S2/H2B. In the competition assays, both full-length and Nap1 ΔA1 interacted with H2A.S2/H2B better than did the Nap1 ΔA3 or Nap1-core domain proteins; Nap1 ΔA3 and Core domain bound histones with relative equal affinity. These results show that the C-terminal A3 acidic stretch increases relative Nap1 affinity for histones.

**Figure 2.**
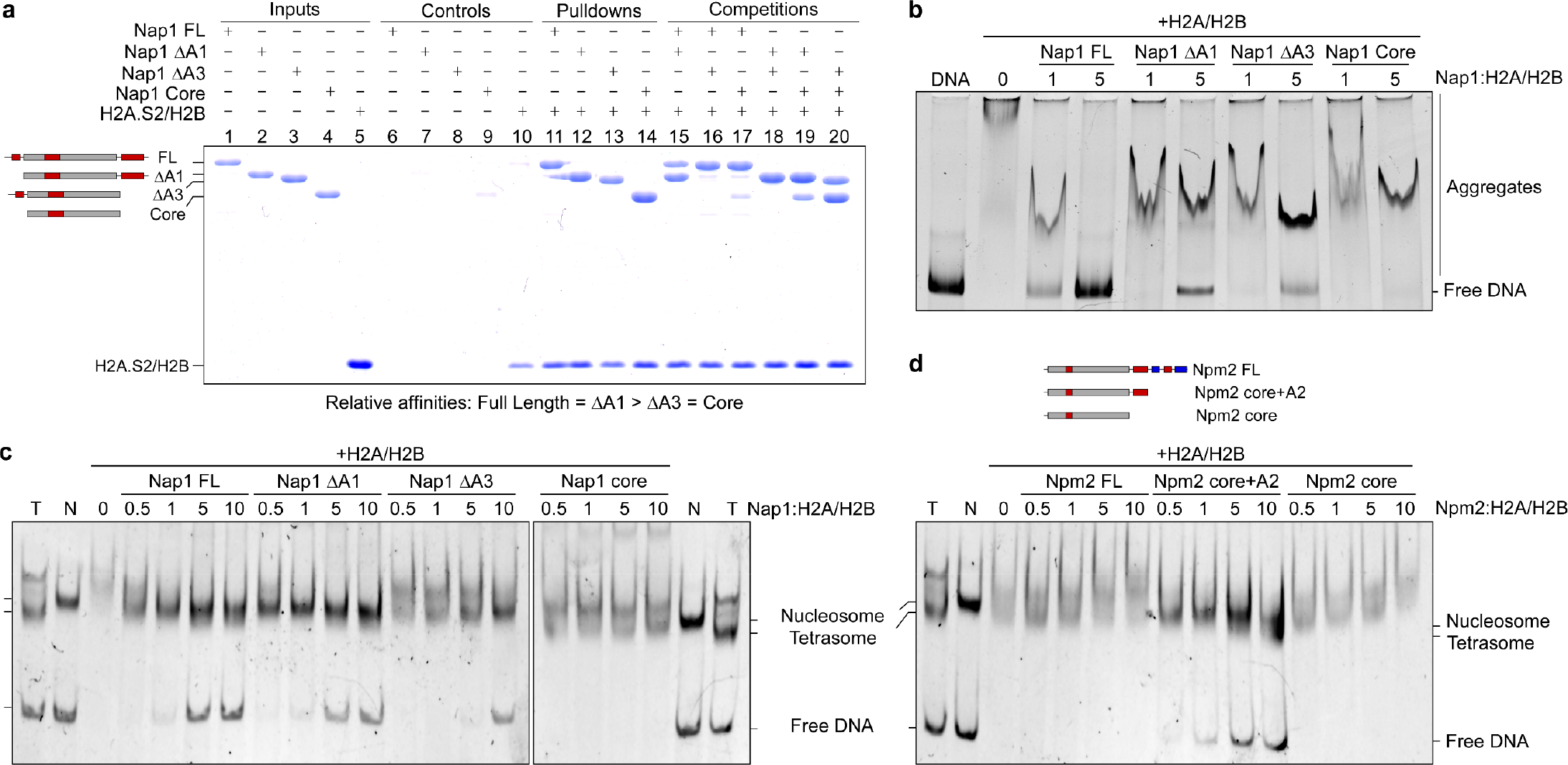
Histone chaperone acidic IDRs maximize histone binding, facilitate disaggregation, and promote nucleosome assembly. **a**. StrepII-tagged H2A.S2/H2B pulldown and competition binding assays of Nap1 truncation mutants. Coomassie stained gel with components included as indicated at the top. Left-graphic shows domain structure of the Nap1 truncations, with gray indicating the core domain and red indicating acidic IDRs. **b**. Histone-capturing disaggregation assay from histone-DNA aggregates using Nap1 truncation mutants. Molar ratio of monomeric chaperone to H2A/H2B is shown **c**. Mononucleosome assembly assay starting from tetrasome (T) using H2A/H2B dimers and Nap1 truncation mutants. Nucleosome (N) included as positive control. The location on the gel of the Nucleosome, Tetrasome, and Free DNA are indicated. molar ratio of monomeric chaperone to H2A/H2B is shown **d**. Same as (c) but using Npm2 truncation mutants.

To assess the role of the acidic tails on the ability of Nap1 to resolve histone-DNA aggregates, we utilized a histone-capturing disaggregation assay (**Figure 2b**). Using a 500bp linear fragment of double-stranded DNA (dsDNA) mixed with a 20-fold molar excess of H2A/H2B heterodimer, we produced a non-nucleosomal histone-DNA aggregate solution. Upon addition to the aggregates, full-length Nap1 captured nearly all histones, resulting in complete recovery of free DNA. The Nap1 ΔA1 construct was partially able to recover free DNA, whereas Nap1 ΔA3 and Core domains were not sufficient to capture histones. These results indicate that both A1 and A3 acidic stretches of Nap1 increase histone binding affinity and are important for its ability to resolve histone-DNA aggregates.

To evaluate the role of the acidic stretches on the ability of Nap1 and Npm2 to form nucleosomes, we utilized a mononucleosome assembly assay (**Figure 2c,d**). Tetrasomes–recombinant H3/H4 assembled with 187 bp of dsDNA by salt dilution–were subsequently mixed with Nap1 or Npm2 prebound to increasing concentrations of H2A/H2B heterodimers; salt-dialysis assembled nucleosomes were included as a positive control. Full-length Nap1 and the ΔA1 construct both formed nucleosomes to a similar extent, whereas ΔA3 did so to a lesser degree; the core domain was incapable of assembling nucleosomes (**Figure 2c**). As we previously showed ^9^, full-length Npm2 contains a C-terminal basic patch that self-associates to block its acidic histone-binding domain. Histone binding is recovered when this basic patch is truncated (Npm2 core+A2). Much like the Nap1 core domain, we previously showed that the Npm2 core domain retains its ability to bind histones. Consistently, only the Npm2 Core+A2 bound to H2A/H2B formed nucleosomes on a tetrasome substrate (**Figure 2d**). Together, these experiments demonstrate that, while the Nap1 and Npm2 acidic tails are dispensable for histone binding, they are required for their histone chaperoning functions of disaggregation and nucleosome assembly.

### Chaperone glutamylation enhances histone binding and promotes histone stabilization

We next sought to investigate the molecular function of post-translational glutamylation of histone chaperones. First, both Npm2 core+A2 and full-length Nap1 proteins were post-translationally glutamylated by recombinant human Tyrosine Tubulin Like Ligase 4 (TTLL4) catalytic domain (**Supplemental Figure S2a**). All TTLL4-treated proteins are hereafter referred to with the suffix “glu”. As we previously showed, recombinant full-length Npm2 is not a substrate for TTLL4 ^30^. Therefore, in these experiments we used the Npm2 core+A2 protein. As we previously showed for Npm2 core+A2 ^9^, the presence of histones H2A/H2B blocked TTLL4 glutamylation of Nap1, consistent with acidic IDR substrates engaging histone binding (**Supplemental Figure S2b**). Glutamylation did not alter the oligomeric state of full-length Nap1 (**Supplemental Figure S2c**).

To test relative interactions, we utilized the H2A.S2/H2B competition pulldown assay with unmodified or glutamylated Npm2 core+A2 and Nap1 proteins (**Figure 3a**). Protein glutamylation resulted in a slight gel retardation (**Figure 3a**, lanes 2 and 4). As a positive control, all four proteins were pulled down by H2A.S2/H2B (**Figure 3a**, lanes 11-14). Unmodified Nap1 had a higher relative affinity compared to Npm2 core+A2 (**Figure 3a**, lane 15). When competed against unmodified proteins, both Npm2 Core+A2glu and Nap1glu have a higher relative affinity for H2A.S2/H2B (**Figure 3a**, lanes 15 and 16). Both glutamylated proteins have similar affinities for H2A.S2/H2B (**Figure 3a**, lane 16). Note that, as Npm2 is a pentamer while Nap1 is likely an oligomer-of-dimers, avidity effects complicate direct affinity comparisons. Overall, these results demonstrate that glutamylation of Npm2 core+A2 and Nap1 increases their relative affinities for H2A.S2/H2B.

**Figure 3.**
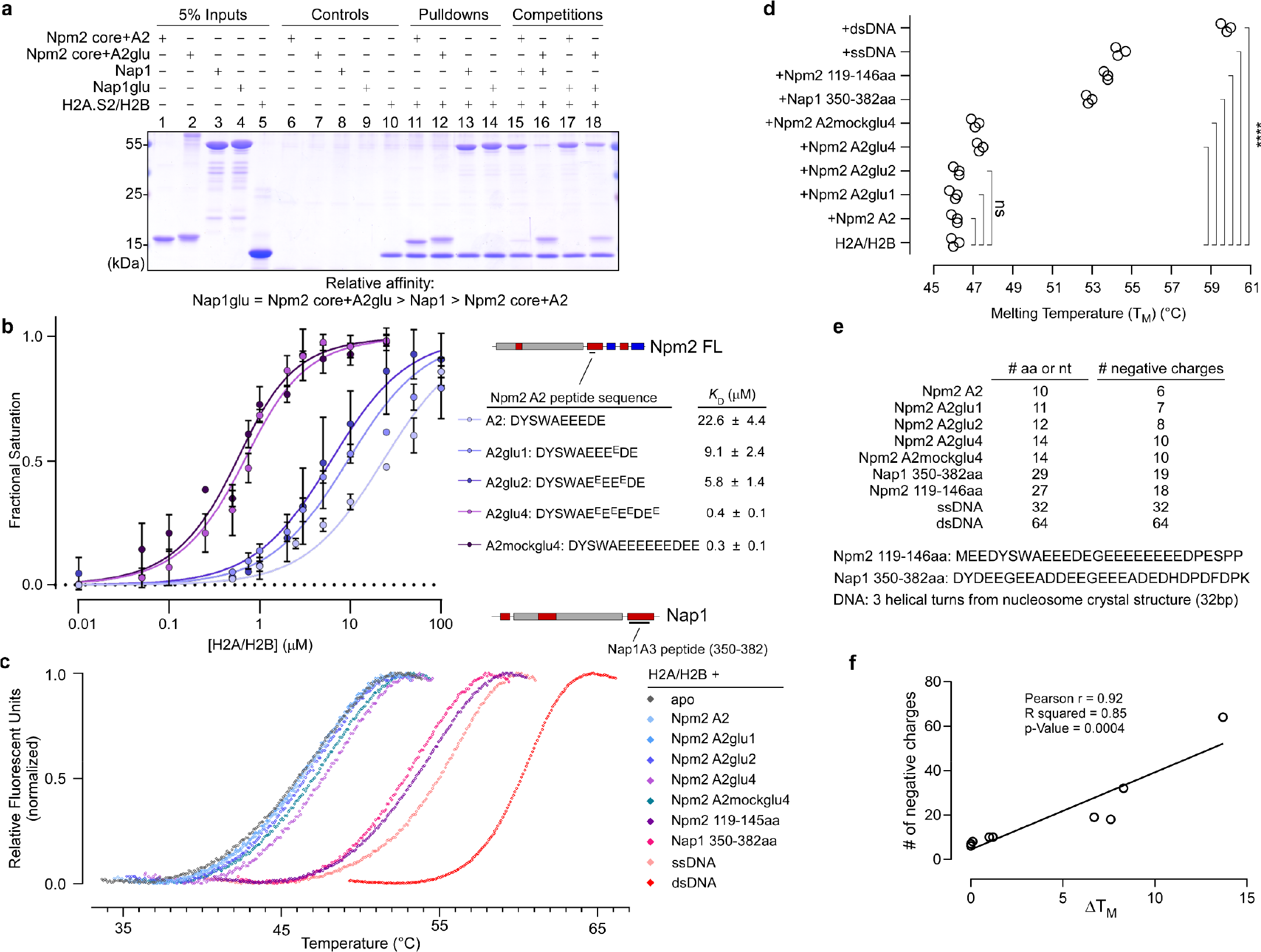
Npm2 and Nap1 acidic IDR glutamylation enhances binding and stabilization of H2A/H2B dimers. **a**. Streptactin H2A.S2/H2B pulldown and competition binding assays of Nap1 and Npm2 core+A2 without or with glutamylation (glu). **b**. Intrinsic tryptophan fluorescence binding assay isotherms using H2A/H2B and Npm2 A2 peptides with increasing glutamylation. Inset: legend and binding constants determined from each isotherm; modified glutamates are indicated with superscript (E^E^). **c**. Normalized thermal shift assay melting isotherms using H2A/H2B dimers and Nap1 or Npm2 peptides or single-stranded or double-stranded DNA **d**. Melting temperatures (T_m_) derived from thermal stability assay isotherms (**** p-val < 0.0001) e. Table of Nap1 and Npm2 peptide amino acid or nucleotide characteristics used in thermal stability assay f. Correlation plot between number of chaperone polyelectrolyte negative charges and change in H2A/H2B melting temperature

We previously demonstrated that, compared to the complete Npm2 Tail, the short Npm2 A2 peptide (^122^DYSWAEEEDE^131^) had relatively weak affinity toward H2A/H2B ^9^. Therefore, we reasoned that glutamylation of this region would substantially increase affinity of Npm2 toward H2A/H2B dimers. To test this, we performed quantitative intrinsic tryptophan fluorescence H2A/H2B binding assays. This assay measures binding-induced environmental changes to the A2 peptide tryptophan. We used the following synthetic unmodified and glutamylated Npm2 A2 peptides: A2glu1-DYSWAEEE^E^DE; A2glu2-DYSWAE^E^EE^E^DE; A2glu4-DYSWAE^E^E^E^E^E^DE^E^; and A2mockglu4 DYSWAEEEEEEDEE, where the superscripted E^E^ denotes sites of glutamate-glutamylation (**Figure 3b**). These experiments showed that, while the unmodified A2 peptide binds to H2A/H2B with modest affinity (*K*_D_ = 22.6 ± 4.4μM), there was a glutamylation-dependent increase in histone affinity: A2glu1 (*K*_D_ = 9.1 ± 2.4μM), A2glu2 (*K*_D_ = 5.8 ± 1.4μM), A2glu4 (*K*_D_ = 0.4 ± 0.1 μM); maximal glutamylation increased the affinity of this peptide toward H2A/H2B by approximately two orders of magnitude.

To separate the effects of specific contacts made due to the branched nature of glutamylation from non-specific electrostatic effects from the extra negative charge on the region, we tested the binding of an A2mockglu4 peptide with 4 extra glutamates added in the primary sequence through normal peptide bonds (DYSWAEEEEEEDEE). We observed that this “pseudo-glutamylated” Npm2 peptide bound to H2A/H2B dimers with high affinity, like that of the glutamylated version (*K*_D_ = 0.3 ± 0.1μM). This result indicated that electrostatic effects may explain the increased affinity observed with the glutamylated peptides.

As histones in solution may be less ordered than when bound to DNA in the nucleosome, we hypothesized that glutamylated acidic IDRs may serve to stabilize the histone fold. To determine if chaperone binding and their post-translational glutamylation increases the thermal stability of H2A/H2B dimers, we performed protein thermal shift melting assays in the absence and presence of various Npm2 and Nap1 acidic peptides; we also tested single-stranded and double-stranded DNA ligands. In the absence of a chaperone peptide, the average melting temperature of recombinant *Xenopus* H2A/H2B dimer was at 46.1 ± 0.2°C (**Figure 3c,d**), similar to that measured previously by differential scanning fluorimetry ^42^. The addition of the short Npm2 A2, A2glu1, and A2glu2 did not change the observed T_m_ of H2A/H2B. The addition of Npm2 A2glu4 and A2mockglu peptides increased the T_m_ of H2A/H2B by 1.2 °C (p-val < 0.0001, one-way ANOVA). The addition of Npm2 Tail (119-146) and Nap1 A3 (350-382) acidic peptides–which are both longer and contain many more aspartate and glutamate residues (**Figure 3e**)–shifted the H2A/H2B T_m_ up by 7.6 °C and 6.7 °C, respectively (**Figure 3d**).

Since glutamylation increases the overall net negative charge of histone chaperone acidic IDRs, we hypothesized that glutamylation may function to increase DNA mimicry. To test this hypothesis, we added single-stranded and double-stranded DNA corresponding to the 32bp sequence of that in the nucleosome core particle crystal structure (PDB: 1KX5) ^38^ to H2A/H2B dimers and performed the thermal shift assay. Single-stranded DNA increased the T_m_ of H2A/H2B by 8.3 °C, like the shift induced by the longer Npm2 and Nap1 peptides. Double-stranded DNA shifted the T_m_ up by 13.7 °C (**Figure 3d**). While the increase in T_m_ of H2A/H2B dimers showed no correlation with overall isoelectric points of the chaperone peptides (**Supplementary Figure S2d**), it correlated well with the number of acidic charges within the peptide and within DNA (**Figure 3f**). Together, these data indicate that binding of long acidic chaperone peptides promotes increased stability of H2A/H2B and that glutamylation of Npm2 and Nap1 acidic stretches functions to further stabilize the heterodimers. Overall, the binding of H2A/H2B to acidic IDRs or DNA exhibited similar changes to thermal stability.

### Glutamylation promotes histone chaperone activity

As we demonstrated that chaperone acidic IDRs and their glutamylation enhances histone binding and stabilization, we hypothesized that it is a potential regulatory PTM for promoting chaperone efficiency. Therefore, our next goal was to investigate how glutamate-glutamylation alters histone chaperone function. To this end, we used both the histone-capturing disaggregation and mononucleosome assembly (**Supplementary Figure S3a**) functional assays. While both unmodified Npm2 core+A2 and Nap1 were able to partially resolve the histone-DNA aggregates, their glutamylated counterparts completely captured histones from the aggregate at a lower chaperone:H2A/H2B ratio (**Figure 4a and 4b**, respectively). Although the Nap1 A1 and A3 acidic patches were dispensable for low-affinity histone binding (**Figure 2a**), we tested the effect glutamylation has on the Nap1 truncations with respect to capturing histones from the DNA aggregate (**Figure 4c**). Glutamylation of Nap1 ΔA1 dramatically increased histone capture in the 1:1 Nap1:H2A/H2B ratio experiment, unmodified ΔA1 modestly recovered free DNA, and glutamylated ΔA1 completely resolved aggregation. The Nap1 ΔA3 truncation showed a similar effect: in the 5:1 Nap1:H2A/H2B ratio experiment, glutamylated Nap1 ΔA3 recovered more free-DNA than did its unmodified counterpart. While short DNA fragments were unable to remove Nap1 or Nap1glu from H2A.S2/H2B (**Supplementary Figure S3b**), these results are generally consistent with acidic tails competing with DNA for histones and with the hypothesis that glutamylation enhances this competition by increasing DNA mimicry.

**Figure 4.**
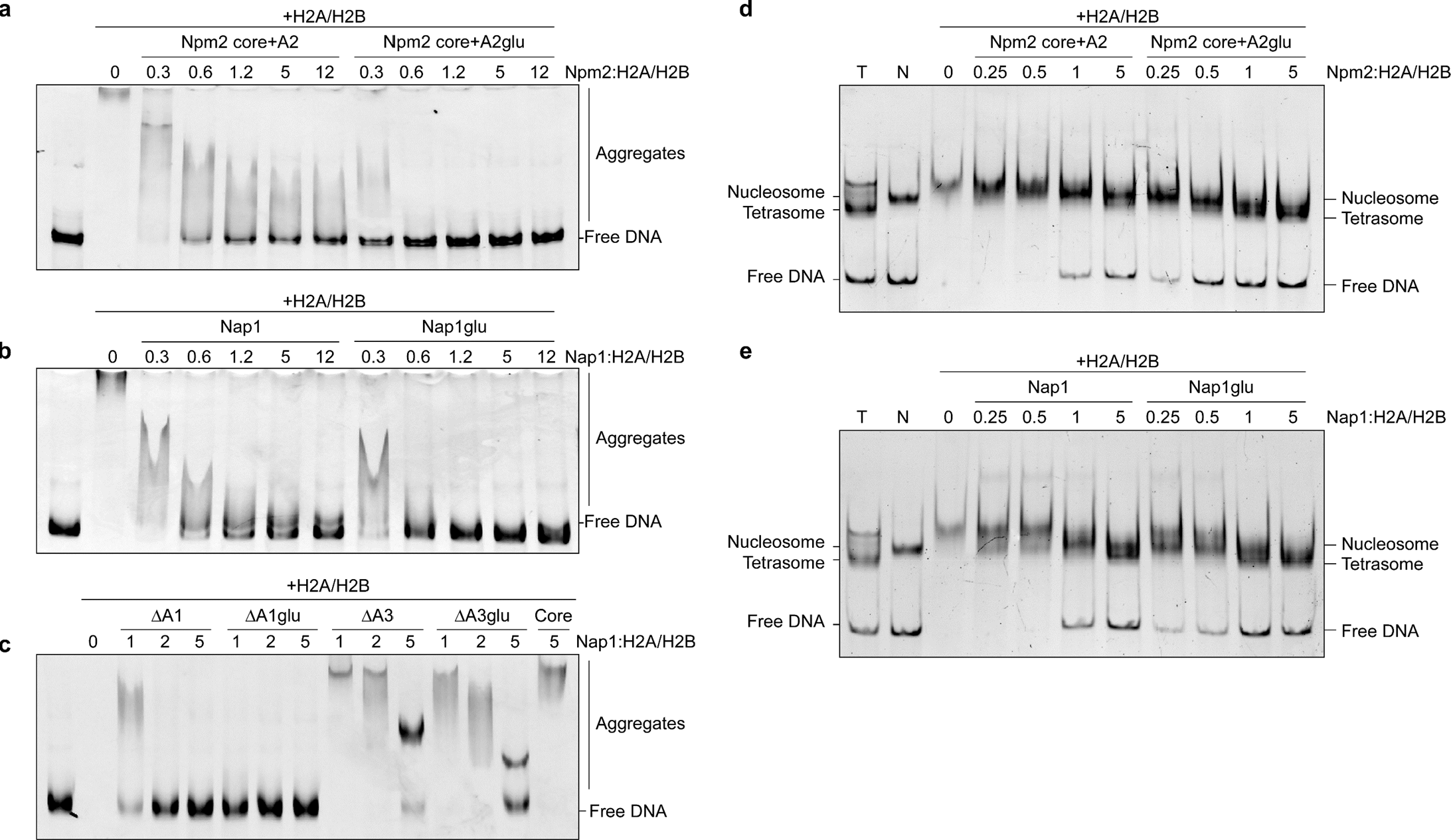
Npm2 and Nap1 glutamylation enhances histone capture and promotes mononucleosome assembly. **a**. Histone-capturing disaggregation assay from histone:DNA aggregates using either unmodified or glutamylated Npm2 core+A2. Chaperones were titrated as indicated, molar ratio of monomeric chaperone to H2A/H2B is shown **b**. Histone-capturing disaggregation assay from histone:DNA aggregates using either unmodified or glutamylated Nap1, molar ratio of monomeric chaperone to H2A/H2B is shown **c**. Histone-capturing disaggregation assay from histone:DNA aggregates using either unmodified or glutamylated Nap1 truncation mutants, molar ratio of monomeric chaperone to H2A/H2B is shown **d**. Mononucleosome assembly assay starting from pre-assembled tetrasomes using H2A/H2B dimers and unmodified Npm2 core+A2 and TTLL4-treated Npm2 core+A2 (Npm2 Core+A2 glu), molar ratio of monomeric chaperone to H2A/H2B is shown **e**. Mononucleosome assembly assay starting from pre-assembled tetrasomes using H2A/H2B dimers and unmodified Nap1 and TTLL4-treated Nap1 (Nap1glu), molar ratio of monomeric chaperone to H2A/H2B is shown

Finally, to measure glutamylation-induced changes in nucleosome assembly activity, we used the tetrasome-deposition assay as described above. In line with the disaggregation assay, glutamylation of Npm2 core+A2 enhanced H2A/H2B deposition on tetrasomes to assembly nucleosomes (**Figure 4d**, compare 0.25 and 0.5 molar ratios in control and TTLL4 treated samples). Similarly, glutamylation of Nap1 enhanced deposition of H2A/H2B on tetrasomes (**Figure 4e**, compare 0.25 and 0.5 molar ratios in control and TTLL4 treated samples); however, neither Nap1 nor Nap1glu were able to disassemble pre-made mononucleosomes (**Supplementary Figure S3c**). Taken together, these results strongly demonstrate that the intrinsically disordered acidic tails of Npm2 and Nap1 are critical for histone chaperoning activity and that glutamylation amplifies their efficiency.

### Molecular dynamics simulations reveal modes of acidic IDR binding to the H2A/H2B heterodimer

To understand how acidic IDRs and glutamylation mechanistically influence chaperone:histone binding and also affect histone protein stabilization, we first used computational approaches. To predict structural changes and contacts that occur upon binding, we performed one microsecond, all-atom, explicit solvent molecular dynamics simulations of the H2A/H2B heterodimer in complex with unmodified or fully-glutamylated Npm2 A2 or A2glu4 (^122^DYSWAEEEDE^131^) or in complex with Nap1 A3 or A3glu9 (^350^DYDEEGEEADDEEGEEEAD-EDHDPDFDPK^382^) peptides; underscored residues correspond to sites of modeled glutamylation. Each trajectory was performed twice, from distinct starting positions; representations are shown from a single simulation (**Supplemental Movies 1a-e**).

As demonstrated in solution with our thermal shift assay (**Figure 3c-f**), binding of the Npm2 A2glu4, Npm2 119-146aa, and Nap1 350-382aa peptides all resulted in significant stabilization of the H2A/H2B heterodimer. To assess how glutamylation of Npm2 A2 and Nap1 350-382aa affected histone stabilization, across the trajectories we measured the average fluctuation of each residue of the H2A/H2B dimer, either in its *apo* form or bound to either unmodified or glutamylated Npm2 A2 or Nap1 peptides (**Figures 5a,b**, respectively). In the simulation, both unmodified peptides stabilized the H2A/H2B dimer—compare apo (blue lines) to unmodified peptides (gray lines)— showing reduced fluctuations across both histone folds. Notable stabilization ranging from the terminal end of the H2A α1 through the α3 helices was observed upon interaction with Npm2 A2 or Nap1 A3 peptides; additional stabilization at the H2A αC helix occurs with Npm2 A2 interaction. Both unmodified Npm2 A2 and Nap1 peptides also reduced fluctuations across the entire H2B fold, ranging from α1 through αC helices. Compared to its unmodified counterpart, the glutamylated Npm2 A2glu4 peptide (**Figure 5a**, orange lines) further reduced fluctuations of H2A, ranging from α2 through α3 helices and had a modest effect on H2B stability. Compared to unmodified Nap1, in the simulation the Nap1 A3glu9 peptide (**Figure 5b**, orange lines) also increased stability of the H2A α3 to αC helices; it also stabilized the 3_10_ helix at the H2A C-terminus. Likewise, in the trajectory Nap1glu9 reduced fluctu-ations in the H2B α2 helix and had modest effect between α3 and αC helices. Overall, *in silico*, glutamylation of Npm2 and Nap1 reduced fluctuations and stabilized both H2A and H2B histone folds.

**Figure 5.**
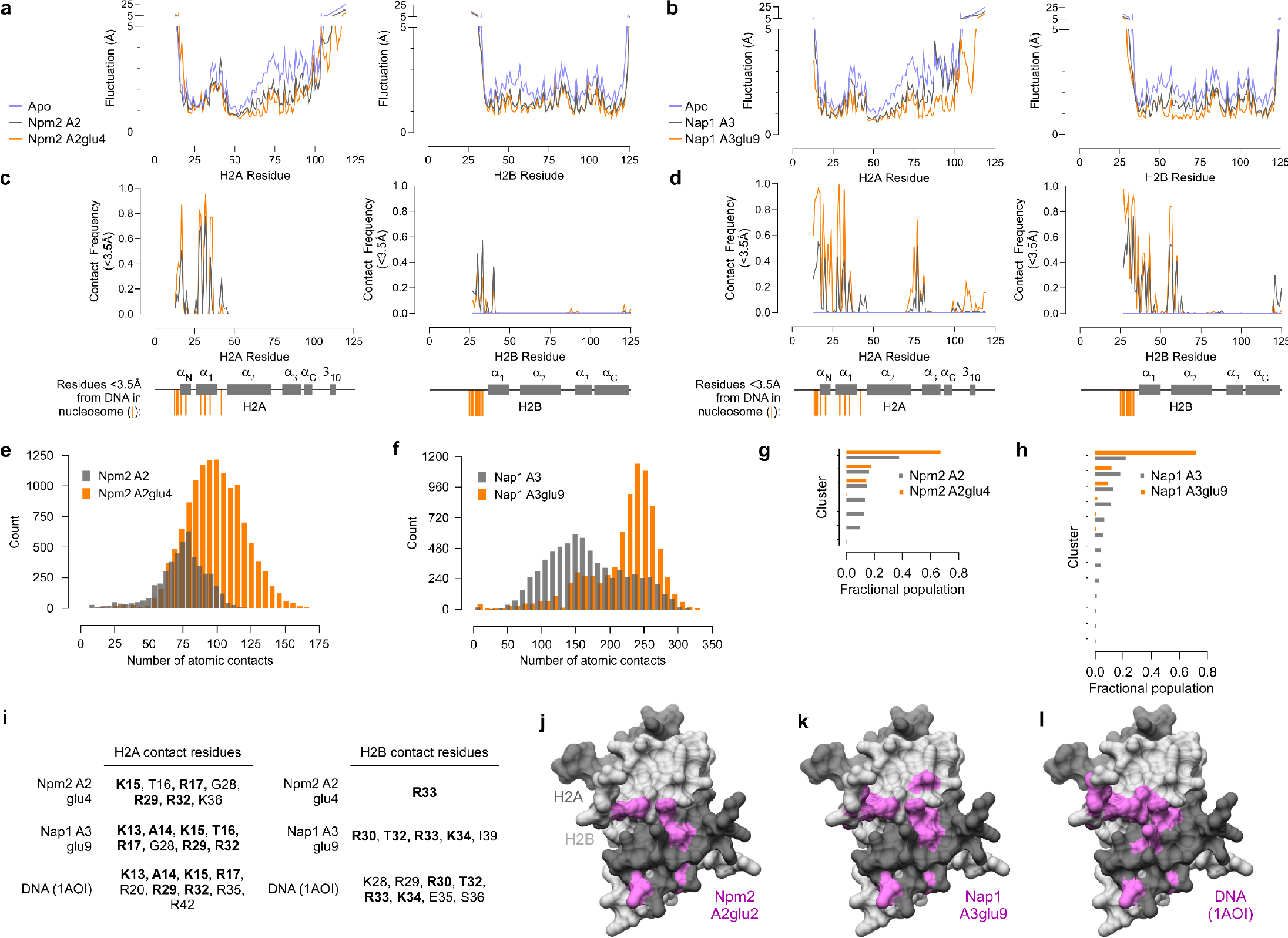
All-atom, 1 μs explicit solvent, molecular dynamics simulations show that Npm2 A2 and Nap1 A3 glutamylation stabilizes histones upon interaction. **a**. Average fluctuation plots of histone residues (H2A (left) and H2B (right)) in the apo (blue), Npm2 A2 (grey) or Npm2 A2glu4 (orange) encounter complexes. **b**. Same as in (a) but for Nap1 A3 (grey) and Nap1 A3glu9 (orange). **c**. Interacting residues between Npm2 A2glu4 or Nap1 A3glu9 and H2A (top) and H2B (bottom); bolded residues are common histone residue contacts between both peptides. Bottom H2A and H2B domain maps, orange vertical lines indicate residues < 3.5Å from DNA in the nucleosome (PDB:1AOI) **d**. Same as in (c) but for Nap1 A3 (grey) or Nap1 A3glu9 (orange) encounter complexes. **e**. Distribution of atomic contacts between H2A/H2B and Npm2 A2 (grey) or Npm2 A2glu4 (orange). **f**. Same as in (g) but for Nap1 A3 (grey) or Nap1 A3glu9 (orange). **g**. Density-based conformational clustering of Npm2 A2 (grey) or Npm2 A2glu4 (orange) conformations when interacting with H2A/H2B. **h**. Same as in (e) but for Nap1 A3 (grey) or Nap1 A3glu9 (orange). **i**. Table of H2A (left) and H2B (right) residues with average distance over the MD trajectories < 3.5Å distance to histone chaperone peptides **j**. Representative models from the most dominant clusters for Npm2 A2glu4 encounter complexes with H2A/H2B. Pink indicates contact residues highlighted in (i) **k**. Same as (j) but for Nap1 A3glu9 **l**. Model of H2A/H2B and DNA from the nucleosome crystal structure (PDB: 1AOI). Pink indicates residues < 3.5Å from DNA

To understand why the glutamylated peptides stabilize the histones, we measured how they interacted. Over the course of the trajectory, the frequency of atomic contacts with the H2A/H2B dimer was both markedly localized and increased by glutamylation of both peptides. We measured the frequency of contact (defined <3.5Å) between each histone amino acid and any chaperone peptide amino acid. As shown in **Figure 5c,d**, glutamylated chaperone peptides (orange) made more frequent contact with localized histone residues than did non-glutamylated peptides (gray); these contact regions were remarkably similar to histone:DNA contacts (<3.5Å) in the nucleosome core particle (**Figure 5c,d**, bottom domain cartoon, orange lines). Across the trajectories, the unmodified Npm2 A2 peptide resulted in a distribution with a mean of 61 contacts (**Figure 5e**, gray bars), whereas the Npm2 A2glu4 peptide shifted this mean to approximately 125 contacts (orange bars). Likewise, the unmodified Nap1 A3 peptide showed a contact distribution with a mean of 166 contacts (**Figure 5f**, gray bars), and the Nap1 A3glu9 peptide shifted the mean to 268 atomic contacts (orange bars).

The effect of glutamylation is also highlighted by differences in the number and population of dominant conformational clusters of the IDR peptides. A density-based conformational clustering analysis showed that the Npm2 A2 peptide forms seven major clusters, with the top populated cluster accounting for 40% of the total population and the next five being similarly populated (**Figure 5g**, gray bars). The A2glu4 had better defined conformations, with only 4 major clusters; the most populated cluster contains almost all the conformations at approximately 70% of the total population (**Figure 5g**, orange bars). Similarly, simulation of the Nap1 A3 peptide binding H2A/H2B resulted in over a dozen sparsely populated clusters, with the top populated cluster accounting for only 20% of the total population (**Figure 5h**, gray bars). The Nap1glu9 peptide had just five clusters, with the top accounting for nearly 75% of the total population (**Figure 5h**, orange bars). These data strongly suggest that glutamylation of these peptides increases both the overall affinity and specificity of histone interactions.

Simulations with the Nap2 A2glu4 or Nap1glu9 peptides showed that Npm2 A2glu4 contacts H2A K15, T16, R17, G28, R29, R32, and K36 and H2B R33. Nap1glu9 contacts H2A K13, A14, K15, T16, R17, G28, R29, and R32 and H2B R30, T32, R33, K34, and I39 (**Figure 5i**). The two glutamylated peptides showed significant overlap with respect to the residues they contact (**Figure 5g**). Despite positioning the peptides randomly and distal from this location at the start of each simulation (**Supplemental Movies 1a-e**), in both simulations both peptides found the same H2A binding pocket early in the MD trajectory (**Figure 5j,k,l**). Supporting the idea that histone chaperone acidic stretches are in direct competition with DNA for histone interaction, this pocket coincides with the residues that directly contact the phosphate backbone of DNA in the nucleosome crystal structure (**Figure 5i**). Together, the results of our MD simulations predict a model in which glutamylation of histone chaperone acidic stretches results in stabilization of H2A/H2B through increased, specific histone residue contacts (**Supplementary Figure S4a-g**).

### Solution NMR assignment and secondary structure of the Xenopus H2A/H2B heterodimer

NMR-studies of the human and *Drosophila* H2A/H2B dimers revealed that, while dynamic in solution, H2A/H2B adopts a similar overall fold as in the nucleosomal counterpart ^7,8^. To test the solution structure and characteristics of *Xenopus* H2A/H2B, we individually labeled either H2A or H2B and formed the dimer with its unlabeled counterpart; these isotopically triple labeled dimers were [U-^13^C,^15^N,^2^H]-H2A/H2B and H2A/[U-^13^C,^15^N,^2^H]-H2B. Using TROSY-style triple-resonance experiments, we assigned 82% of possible Cα, Cβ, CO, HN, and N chemical shift values for H2A, and 74% of possible Cα, Cβ, CO, HN, and N chemical shift values for H2B. The unstructured tails of both H2A/H2B were challenging to assign; overall, we assigned 93% of H2A and 85% of H2B residues from the folded histone core in the HSQC-spectra (**Figures 6a and 6b**, respectively).

**Figure 6.**
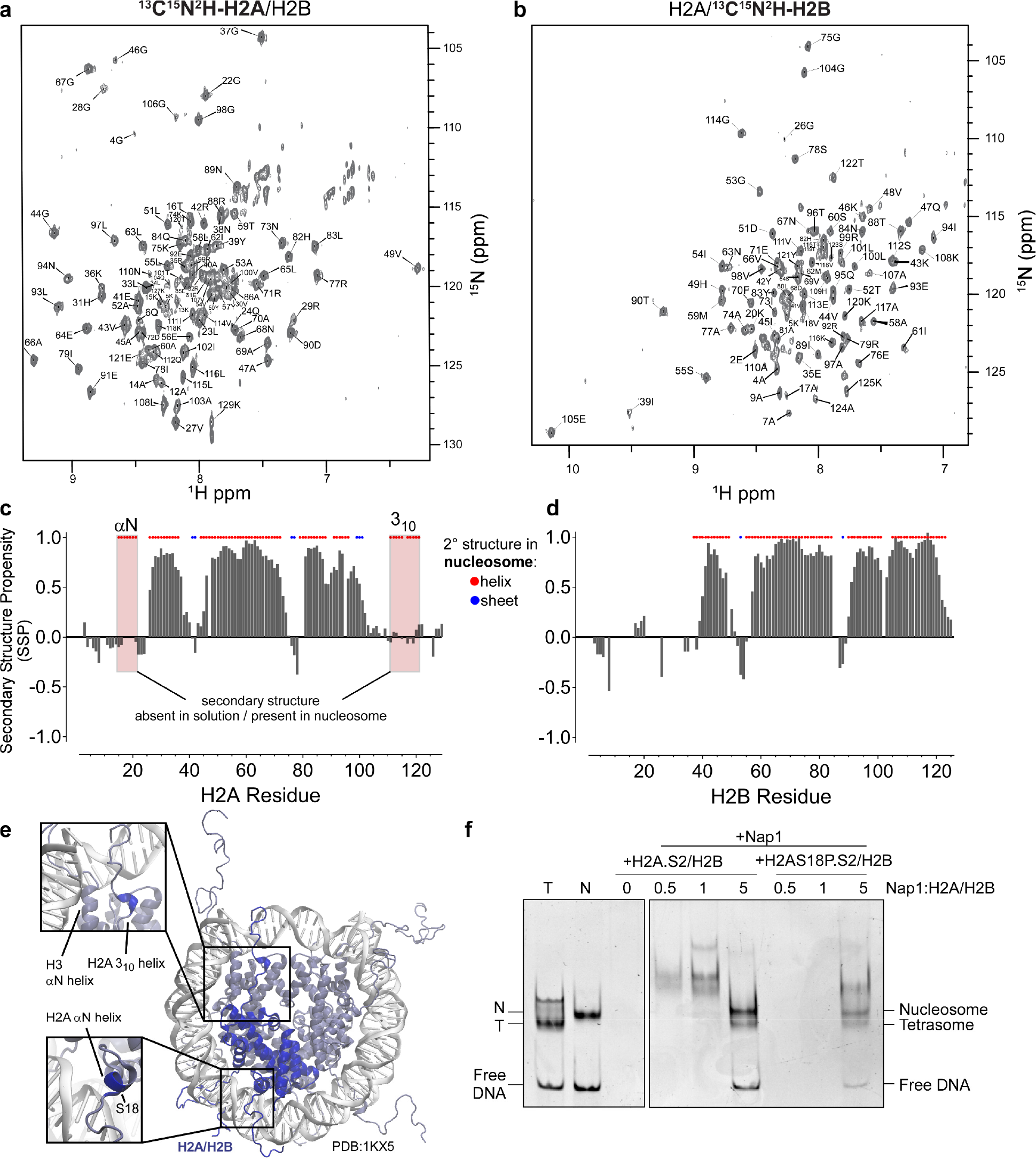
Solution NMR analysis of the *Xenopus laevis* H2A/H2B dimer reveals secondary structure differences with the nucleosome. **a**. Assigned ^1^H-^15^N-TROSY-NMR spectra of triple labeled ^2^H-^13^C-^15^N-H2A in complex with H2B. **b**. Assigned ^1^H-^15^N-TROSY-NMR spectra of triple labeled ^2^H-^13^C-^15^N-H2B in complex with H2A. **c**. Secondary structure propensity (SSP) of H2A (histogram) compared with secondary structure of H2A in nucleosome (red dots = helical residues; blue dots = beta sheet). **d**. Same as (C) but for H2B **e**. Representation of nucleosome core particle (PDB:1KX5) with H2A/H2B chains C,D shown in blue (center). Bottom inset shows the H2A αN helix adjacent to DNA. Top inset shows H2A C-terminal 3_10_ helix adjacent to the H3 αN helix. **f**. Mononucleosome assembly assay starting from pre-assembled tetrasomes using wildtype H2A.S2/H2B (left) or S18P mutant (right) dimers and unmodified Nap1. N (Nucleosome), T (Tetrasome), and Free DNA locations are indicated.

To ascertain the stability of the individually labelled dimers, we also performed variable-temperature (VT) NMR experiments. We recorded a series of ^1^H-^15^N HSQC spectra of the unbound [U-^2^H,^15^N]-H2A/H2B and H2A/[U-^2^H,^15^N]-H2B dimers as a function of temperature (40°C, 25°C, 16°C, 10°C and 4°C). Temperature-induced changes in protein dynamics and from conformational exchange processes result in line broadening and chemical shifts in NMR spectra. Lowering the temperature of these samples showed near linear decreases in peak intensities for nearly all residues within the histone fold, while peaks corresponding to residues of the histone tails remained intense at lower temperatures (**Supplemental Figure S5a,b**). In contrast, when raised to 40°C, peak broadening and CSP occurred globally. This could be attributed to the temperature induced increase in the thermal motion of atoms, which causes more mobility and thus line broadening effect, while CSPs appear due to changes in the chemical environment due to increased dynamicity. In both cases, returning the samples back to room temperature led to re-appearance of the peaks corresponding to residues in the histone fold, with intensities comparable to the samples prior to heating or cooling (*not shown*). These results indicate that the temperature-induced peak loss observed is reversible and likely caused by changes in dynamics of the H2A/H2B dimer and not by irreversible protein aggregation. Therefore, all further experiments were conducted at 25 °C.

To determine if in solution H2A/H2B folds into a similar conformation as does nucleosomal H2A/H2B, we performed secondary structure propensity (SSP) analysis on assigned residues from the *Xenopus* H2A/H2B dimer ^43^. Comparing the SSP values to secondary structure elements found in a high-resolution crystal structure of the nucleosome (PDB 1KX5) ^44^ showed that the H2A/H2B dimer similarly folds in solution (**Figure 6c and 6d**, respectively). The most noticeable differences are in H2A, in which the αN and two C-terminal 3_10_ helices are found in the nucleosome crystal structure but are not detected in the H2A/H2B dimer in solution ^38,44^. These elements are absent in the NMR structure of the human and *Drosophila* H2A/H2B dimer ^7,8^. Furthermore, both the αN helix and two short C-terminal 3_10_ helices of H2A make direct contacts with the phosphate backbone of DNA in the structures of the nucleosome, suggesting that these helices may form either upon chaperone binding or during nucleosome formation for proper incorporation of H2A/H2B (**Figure 6e**). We also noted that the αC of H2A appears extended in solution compared to the nucleosome structure. Unlike the αN and 3_10_ helices, the αC helix of H2A is buried in the middle of the nucleosome structure and makes no direct contact with DNA. However, it makes multiple direct contacts with the α3 and αC helices of H2B in the nucleosome structure, and extension of this helix would likely cause steric clashes with the C-termini of both H2B and H4 in the nucleosome structure. These results indicate that, while *Xenopus* H2A/H2B dimers in solution largely resemble nucleosomal H2A/H2B, they exhibit structural features that must change prior to nucleosome incorporation. To test if αN formation is necessary for chaperone-dependent nucleosome assembly, we compared Nap1 deposition of wildtype or H2AS18P/H2B; this S18P mutant is predicted to be helix destabilizing. Supporting the hypothesis that pre-formation of this helix is important, we observed reduced deposition of this mutant dimer (**Figure 6f**).

### Chemical Shift Perturbation (CSP) Mapping of Npm2 peptide binding to the H2A/H2B dimer

To determine consequences of chaperone acidic IDR interactions with histones in solution, we utilized chemical shift perturbation (CSP) and peak broadening (peak intensity) analysis of chaperones titrated against either [U-^2^H,^15^N]-H2A/H2B or H2A/[U-^2^H,^15^N]-H2B samples. As our prior analysis revealed that the Npm2 C-terminal Tail domain (residues 119-146) tightly bound H2A/H2B ^9^, we first determined gross changes upon Npm2 Tail binding. Upon titration of the Tail domain the most evident change was the global loss of peak intensities for nearly all residues within the histone fold (**Supplemental Figure S5c**). Decreased peak intensity can arise from exchange processes, conformational exchange, or rapid binding/unbinding dynamics. Surprisingly, most of these residues (25-100 for H2A, and 39-123 for H2B) decreased to ∼50% of their original intensity after binding, whereas peaks corresponding to the disordered histone tails remained intense in both the bound and unbound state.

To delineate the specific histone interactions of the acidic IDRs of Npm2, we performed similar CSP experiments using the short Npm2 A2 peptide (^122^DYSWAEEEDE^131^). Upon titration of this peptide into labeled histones, we observed that peaks remained at a similar intensity as in the unbound state, with most residues in the histone fold only decreasing to ∼75% of their original intensities (**Supplemental Figure S5c**, second row). CSPs were significantly smaller than with the complete Tail domain, with very few residues shifting >0.05ppm. All residues that shifted significantly upon titration of this peptide also shifted in the Tail titration experiments, indicating a similar binding site. For the [U-^2^H,^15^N]-H2A/H2B sample, significant CSPs (>0.05ppm) were observed for residues V27, G28, R29, and V100 (**Figure 7a**). For the H2A/[U-^2^H,^15^N]-H2B sample significant CSPs (>0.05ppm) were observed for residues I39, T119, T122 (**Figure 7b**). Compared to the Tail titration, there appeared to be a greater preference for the C-terminus of H2A and the N-terminus of H2B with this short peptide, possibly indicating weaker binding site specificity.

**Figure 7.**
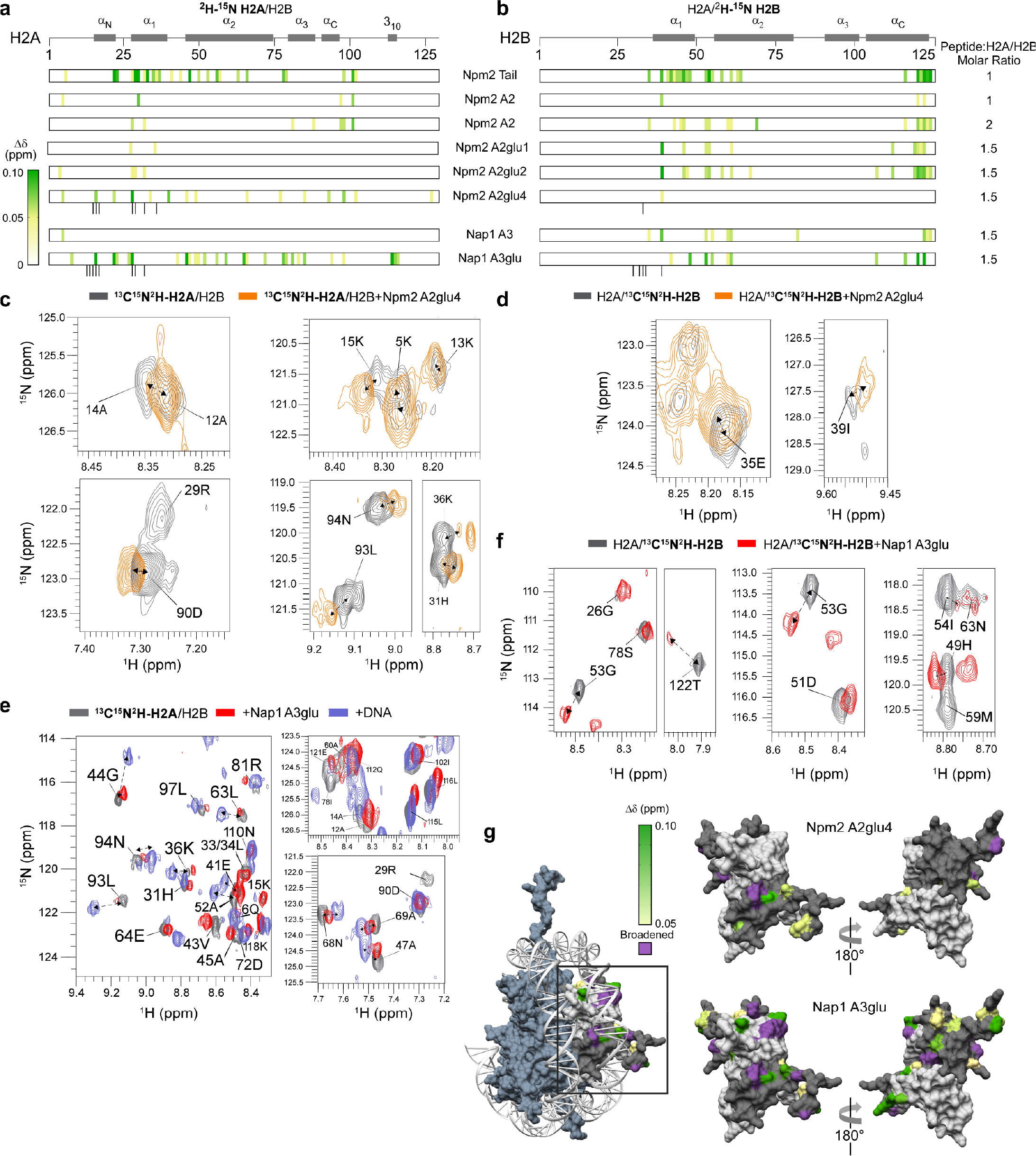
Glutamylated acidic IDRs bind H2A/H2B’s DNA interacting surface. **a**. Heatmap representation of indicated histone chaperone peptide induced chemical shift perturbations (CSP, scale as indicated on left index, yellow to red) across H2A assigned residues in dimer with H2B. Domain structure of H2A is shown. **b**. Heatmap representation of indicated histone chaperone peptide induced chemical shift perturbations (CSP, scale as indicated on left index, yellow to red) across H2B assigned residues in dimer with H2A. Domain structure of H2B is shown. **c**. ^2^H-^15^N-H2A residues (gray) undergoing CSP and/or line broadening upon Npm2 A2glu4 peptide addition (1.5X molar ratio peptide:histone, orange). **d**. ^2^H-^15^N-H2B residues (gray) undergoing CSP and/or line broadening upon Npm2 A2glu4 peptide addition (1.5X molar ratio peptide:histone, orange). **e**. ^2^H-^15^N-H2A residues (gray) undergoing CSP and/or line broadening upon either Nap1 A3glu peptide or DNA addition (1.5X molar ratio peptide:histone, red; DNA:histone, blue). **f**. ^2^H-^15^N-H2B residues (gray) undergoing CSP and/or line broadening upon Nap1 A3glu peptide addition (1.5X molar ratio peptide:histone, orange). **g**. Histone H2A (dark gray) and H2B (light gray) from the nucleosome structure shown, with CSPs (green scale) and broadened (purple) residues highlighted for both Npm2A2glu4 (top) and Nap1 A3glu (bottom) peptide.

Next, to test molecular consequences of glutamylation on histone binding, we used glutamylated synthetic unmodified and glutamylated Npm2 A2 peptides. Addition of the synthetic glutamylated peptides (A2glu1, A2glu or A2glu4) to the labelled dimers exhibited a concentration dependent increase in peak broadening and CSP for almost all residues (**Supplemental Figure S5d**). Such profound effect on the overall dynamics and conformation of the histone dimer is consistent with the stronger binding affinity of the glutamylated peptides compared to the unmodified peptide. Of note, residues in the histone fold that appeared to be actively involved in the interaction with the longer tail peptide depicted a higher degree of perturbation upon addition of the glutamylated Npm2 A2 glu4 peptide. Additionally, residues in the H2A N-terminal helices A12, A14, K13, K15, T16, G28, R29, H31, and R32 in [U-^2^H,^15^N]-H2A/H2B, also predicted by MD simulation studies to be important for interaction undergo significant changes (**Figure 7c**). Ligand binding often leads to conformational changes in the protein that propagates to distant regions, altering the chemical environments of nuclei that are not directly involved in binding. In line with this hypothesis, we also observed a few residues in the C-terminal helix of [U-^2^H,^15^N]-H2A/H2B namely, D90, E92, L93, and N94 depicting similar changes. In the case of H2A/[U-^2^H,^15^N]-H2B, residues I39, S123, A124, and K125 were additionally affected (**Figure 7d**).

### Chemical Shift Perturbation (CSP) Mapping of Nap1 peptide binding to the H2A/H2B dimer

Next, we sought to do similar binding experiments using the longer Nap1 peptide. Compared to the Npm2 tail peptide, the Nap1 peptide interacted much more strongly with the histone dimers, with almost up to 50% peak broadening for majority of the residues in both [U-^2^H,^15^N]-H2A/H2B and H2A/[U-^2^H,^15^N]-H2B when compared to the unbound state (**Supplemental Figure S6d**). Consistent with the MD simulations, this observation could be attributed to the longer peptide length providing a higher surface area of interaction. Like the Npm2 A4glu peptide interacting sites, most of these residues belonged to the histone fold region. Certain others included residues, R36, R77, L83, S113, and L116 in [U-^2^H,^15^N]-H2A/H2B and G26, E35, K46, Q47, A58, M59, S64, and residues in the C-terminal 3_10_ helix K120, and Y121 as predicted by the MD studies (**Figure 7e**). Compared to this, the glutamylated variant of Nap1 peptide showed greater CSP for almost all the residues in the histone fold regions. Residues like T16, H31, L65, L93, and N94 in [U-^2^H,^15^N]-H2A/H2B and I39, T122, S123, and K125 in H2A/[U-^2^H,^15^N]-H2B showed higher CSP and/or broadening when compared to non-glutamylated variant (**Figure 7f**). Finally, these conserved H2A/H2B DNA binding, chaperone peptide interacting regions were also observed on histone peptide array studies with full-length Npm2 ^9^ and full-length Nap1 (**Supplemental Figure S6e**). Overall, these observations support chaperone IDR specific and localized H2A/H2B interactions on DNA binding surfaces, greatly amplified by the presence of glutamate glutamylation.

## DISCUSSION

Due to the lack of structural conservation, generalized histone chaperone mechanisms have long been elusive. In this work, we tested and validated the hypothesis that histone chaperone acidic IDRs act as DNA mimetics to pre-stabilize histones H2A/H2B for deposition and also to remove DNA:histone aggregates. Using biochemical analysis of the canonical−but structurally and biologically unique−*Xenopus laevis* histone chaperones Npm2 and Nap1, we directly show the mechanistic function of these acidic tails. We confirmed glutamate-glutamylation of these acidic tails enhance their function in histone binding, histone:DNA aggregate resolution, and nucleosome assembly (**Figure 7g**). MD simulations and solution NMR experiments supported our biochemical findings by revealing intricate interactions that lead to increased DNA mimicry. The simulations demonstrate that glutamylation of these acidic IDRs promotes contact with H2A/H2B, resulting in increased histone fold stability. Biophysically testing these binding predictions, NMR studies revealed distinct secondary structural differences in H2A/H2B, suggesting that these chaperone peptides might interact more favorably in a nucleosome-like conformation. The NMR data show that the glutamylated regions interact at the same sites as DNA does on the histones, demonstrating that these modifications function to increase DNA mimicry. This discovery significantly broadens our understanding of post-translational modifications in mediating stabilization of H2A/H2B heterodimers and promoting chaperone efficiency.

Our NMR assignment and secondary structure analysis of the *Xenopus* H2A/H2B dimer shows that while most of the histone fold is identical to that observed in the nucleosome structure, the short N-terminal helix of H2A does not form in solution. This finding is also consistent with the NMR structural analysis of the human H2A/H2B dimer in solution ^7^. This helix is present in every published nucleosome crystal structure and makes direct contact with the phosphate backbone of the DNA. This likely indicates that this helix must form for H2A/H2B to be properly deposited onto DNA to form nucleosomes. Chaperone binding may induce the formation of this helix, thus promoting the deposition of H2A/H2B onto DNA. Supporting this hypothesis, we previously detected contacts between the Npm2 Tail domain and a peptide corresponding to this N-terminal helix in peptide array experiments, and our previous PRE-NMR structural models position A2 directly adjacent to this region ^9^. H2A S18P mutant histone dimer exhibited reduced Nap1-dependent deposition on pre-existing tetramers, consistent with formation of this helix as an essential step. We detected large CSPs adjacent to this region upon titrating both Npm2 and Nap1 peptides. Due to peak broadening and unassigned residues, we were unable to determine if this helix forms upon chaperone binding; however, these CSPs could arise via direct binding, binding induced structural changes, such as helix formation, or a combination of the two. As our MD simulations started with the nucleosome core particle structure and were of likely too short a time scale to observe helix formation, atomic resolution structures of the H2A/H2B dimer bound to Npm2 and Nap1 acidic peptides may be required to determine if this helix is present in the complex. To that end, our crystallization trials with both H2A/H2B full length dimers and our single-chain H2B-H2A crystal form ^45^ with chaperone peptides were unsuccessful.

Overall, we propose that the acidic IDRs act as dynamic scaffolds for transient, yet high-affinity interactions with histones H2A/H2B. Indeed, in our MD simulations, both the apo and glutamylated peptides approach the histone H2A/H2B surface, within ∼50 ns. Highlighting its enhanced interaction and stability with the histone dimer compared to the apo peptide, the glutamylated peptide forms a more stable and specific ensemble over an extended timescale. Our observations are similar to the recent studies of extreme protein disorder in the prothymosin α linker histone chaperone, which revealed its behavior as a protein polyelectrolyte ^39,46^. Biological polyelectrolyte domains, modulated through the glutamylation of acidic disordered regions, may therefore have a unique ability to maintain highly responsive chaperone regulatory networks by altering histone affinity. This suggests that the glutamylation-driven enhancement of a polyelectrolyte state serves as a mechanism to tune the responsiveness of cellular processes. Glutamate-glutamylation may therefore enable adaptation to changing environmental signals or conditions and may be pivotal for the robustness and efficiency of cellular networks. These findings highlight the multifaceted roles of post-translational modifications and underscore the importance of further studies to explore their regulatory potential.

## LIMITATIONS OF THIS STUDY

A limitation of this study is that we did not test glutamylation consequences in a biological context. While testing histone chaperone glutamylation in cells is of critical importance to understanding the biological ramifications of this modification, it is experimentally challenging in that: core histone chaperones exhibit significant redundancy, making consequences poorly testable; 2) multiple and dense substrate glutamates (e.g. nine in the Nap1 A3 alone) make mutagenesis and outcomes complicated; 3) TTLL4 has multiple cellular substrates, making knock-down consequences unclear. This study was also primarily limited to histone H2A/H2B, and the role of chaperone acidic IDRs and glutamylation on H3/H4 and linker histones are unknown. Future studies on histone glutamylation in a biological consequence would be likely well-served by studying histone variant or linker histone deposition. NMR studies were limited to chemical shift perturbation and intensity measurements; while additional experiments could expand our quantitation of exchange details, they would not impact conclusions regarding glutamylated chaperone IDR DNA mimicry. Finally, while glutamylation has also been observed in mammalian histone chaperones, these studies were only performed in *Xenopus* Npm2 and Nap1.

## EXPERIMENTAL METHODS

### Chemical Reagents, Antibodies, and Peptides

All chemical reagents were procured from Thermo Fisher Scientific (Waltham, MA), Sigma-Aldrich (St. Louis, MO), Research Products International (Mount Prospect, IL), Cambridge Isotopes Laboratory (Andover, MA), IBA Lifesciences (Göttingen, Germany), or Gold Biotechnology (Olivette, MO). The antibodies against *Xenopus laevis* Npm2 and Npm2 core domain were generated by Lampire Biological Laboratories (Pipersville, PA), using recombinant Npm2 full-length and core domain truncation antigens, respectively. The antibody against *Xenopus laevis* Nap1 was generated by Lampire Biological Laboratories (Pipersville, PA), using recombinant Nap1 antigen. The anti-monoglutamylation antibody TTβIIIglu (referred to here as anti-Glu) was generously provided by Dr. Anthony Spano and Dr. Anthony Frankfurter ^47^. Synthetic peptides corresponding to Npm2 A2 and glutamylated counterparts were obtained from GenScript (Piscataway, NJ). The Npm2 119-146aa peptide was purified as previously described ^9^. The cDNA sequence corresponding to Nap1 350-382aa peptide containing the A3 acidic patch was cloned into the pRUTH5-GST vector which produced the peptide fused to a TEV-cleavable N-terminal His_6_+GST tag; expression and purification of the fused peptide is as described below for Nap1 proteins. Following cleavage of the peptide, the His_6_+GST tag and TEV protease were removed by subtractive Ni-NTA chromatography. The Nap1 peptide was lyophilized and further purified by reverse-phase high-performance liquid chromatography using a 250 × 10 mm 4mm Synergi Fusion-RP 80Å C12 column (Phenomenex, Torrance, CA), as previously described ^48^.

### Molecular modeling and pK_a_ prediction of glutamate and γ-glutamyl-glutamate

Maestro (Shrödinger) was used to model electrostatic potential surface maps and predict p*K*_a_ values for glutamate and γ-glutamyl-glutamate. First a glutamate molecule was modeled using the software’s “Build” feature by selecting Add Fragments > Amino acids > Glu. To model γ-glutamyl-glutamate, another glutamate was modeled and the backbone NH_2_ group was deleted; the terminal OH group of the initial glutamate was changed to an NH_2_ group then this NH_2_ group and the Cα of the second glutamate were highlighted; right-clicking and selecting “Add bond” joined the molecules. Molecules were relaxed by clicking Edit > Minimize > All atoms. The “Poisson-Blotzmann ESP” task was used to generate electronic maps of the unmodified and modified sidechains. Electrostatic potentials are displayed from −10 to 10 *k*_B_T/e [*k*_B_, Boltzmann constant; T, temperature (kelvin); e, electron charge]. Surface transparencies were set to 30% front and 10% back from the “Surface display options” menu. p*K*_a_ values were predicted using Epik by selecting the “Emprical p*K*_a_” task.

### Cloning, Expression, and Purification of recombinant proteins

All histones used in this study were purified and refolded as previously described ^49^. Npm2 proteins were purified as previously described ^30^. cDNA sequences corresponding to *Xenopus laevis* Nap1L.1L isoform X2 (Nap1) constructs used in this study were cloned into pRUTH5 expression vectors using ligation independent cloning ^50^, which resulted in fusion proteins containing an N-terminal His_6_ affinity tag followed by a Tobacco Etch Virus (TEV) protease site. The cDNA sequence corresponding to the catalytic domain of human Tubulin Tyrosine Like Ligase 4 (TTLL4, 561-1199aa) was similarly cloned in to pRUTH5-GST expression vector, resulting in a protein fused to a cleavable N-terminal His_6_+GST tag. TTLL4 catalytic domain was only soluble if the His_6_+GST tag remained fused to the protein. Npm2 and Nap1 proteins were expressed in Rosetta2 (DE3) *E. coli* grown in LB medium containing appropriate antibiotics to OD600 ∼0.7 at 37 °C with agitation before induction with 0.5mM IPTG followed by 18 h incubation at 30 °C with agitation. Cells were pelleted by centrifugation for 20 min at 4000xg and resuspended at a 1:10 mass (g) to volume (mL) in lysis buffer (50 mM Tris-HCl pH 8.0, 1 M NaCl, 5 mM *β*-ME, 5 mM imidazole, 10% glycerol). Lysis was performed using an Emulsiflex C3 homogenizer (Avestin) at 4 °C with two passages at approximately 12,000psi. The lysate was centrifuged at 14,000 x g for 45 min and the soluble fraction was incubated with 1 mL HisPur Ni-NTA resin (Thermo) per 1 L of cell culture at 4 °C with agitation for 1 h. Ni-NTA gravity-flow columns were packed in disposable chromatography columns (Bio-Rad) and washed extensively with 25-50 column volumes (CVs) of lysis buffer, followed by 25CVs of the same buffer containing 30mM imidazole; proteins were eluted with 3CVs of the same buffer containing 300mM imidazole. Eluate containing Npm2 or Nap1 protein was transferred to dialysis tubing (10,000 MWCO) and the N-terminal His_6_ affinity tag was cleaved by adding 6x-histidine-tobacco etch virus (TEV) protease catalytic domain at a 1:50 mass ratio (TEV:Nap1) then dialyzed for 16 h at 4 °C against buffer containing 50 mM Tris-HCl pH 8.0, 150 mM NaCl, 5 mM *β*-ME, and 10% glycerol. Subtractive Ni-NTA chromatography was then performed to remove the affinity tag and TEV protease. Proteins were concentrated to ∼15 mg/mL and polished by size exclusion chromatography (Superose 6 Increase, GE) using the aforementioned buffer. Fractions containing pure protein were pooled and concentrated to ∼15 mg/mL, aliquoted, and stored at −80 °C until needed. For GST-TTLL4, expression and purification were carried out in a similar manner, but Rosetta 2 cells were grown in TB medium with appropriate antibiotics.

### Purification of endogenous Nap1 from Xenopus egg extract

Frogs were handled according to IACUC-approved protocols. *Xenopus laevis* egg extract and low speed supernatant (LSS) were prepared as previously described ^51^. Endogenous egg Nap1 (eNap1) was purified (see schematic, **Supplemental Figure S1a**) from 10 mL of LSS supplemented with Neutravidin to a final concentration of 10 mg/mL. Histones H2A.S2 (bearing a C-terminal StrepII-tag, IBA Lifesciences) and H2B were expressed individually, purified under denaturing conditions, and refolded into dimers as described previously^49^. H2A.S2/H2B was added to Streptactin resin (IBA Lifesciences) pre-equilibrated in egg lysis buffer (ELB: 10mM HEPES-KOH pH 7.8, 50 mM KCl, 2.5 mM MgCl_2_) at ratio of 0.4 mg H2A.S2/H2B dimers per 1 mL Streptactin resin and incubated with rotation for 60 min at 4 °C. The resin was then washed 3 x with 5 mL ELB and added to LSS at a ratio of 1μg H2A.S2/H2B dimers per 0.02 mL LSS and incubated with rotation for 90 min at 4 °C. The resin was washed 2 × 5 mL in ELB supplemented stepwise with 125 mM, 250 mM, 500 mM, 1 M, and 2 M NaCl. Both Nap1 and Npm2 were detected in the 0.5 M and 1 M NaCl fractions which were then combined. Npm2 was subsequently immunodepleted using Npm2 antibodies, as verified by immunoblot. eNap1 was buffer exchanged into 50 mM Tris-HCl pH 8.0, 25 mM NaCl, 5 mM *β*-ME and polished by size exclusion chromatography (Superdex 200 Increase, GE). Fractions containing pure eNap1 were pooled and concentrated to ∼2 mg/mL (as measured by SYPRO Orange stain), aliquoted, and stored at -80 °C until needed. eNap1 was confirmed by immunoblot using anti-Nap1 and anti-Glu antibodies.

### Mass spectrometry

Digestion buffer was prepared with 100 mM ammonium bicarbonate, occasionally including 2 M urea buffer to mildly denature the protein, and the pH was adjusted to approximately 7.5 with 10% acetic acid. The samples of endogenous egg Nap1 or recombinant Nap1 were diluted 1:1 with digestion buffer. To reduce the disulphide bonds, dithiothreitol was added to the sample to a concentration of 2 mM and heated to 50 °C for 30 min. Iodoacetamide was prepared and added to a concentration of 6 mM and incubated in the dark for 30 min. Endoproteinase Arg-C (cleaves c-terminal to arginine) or Trypsin (cleaves c-terminal to lysine/arginine) was added in a 1:100 (w/w) ratio with Nap1. Samples were incubated either at 25 °C for 16 h or at 37 °C for 6 h. Samples were frozen until they were evaluated by LC-MS/MS.

The samples were analyzed using in-house prepared columns. Both analytical and pre-columns were made of fused silica (360 μm OD x 75 um ID for analytical and 100 um ID for pre-column) and had a 2 mm kasil frit. Analytical columns had a laser pulled tip. Two separate packing materials were used to evaluate samples: shorter peptides were evaluated with Dr. Maisch C18 packing material, both 3 and 5 μm particle size, packed to 10 cm; larger peptides were evaluated with Agilent Poroshell, SB-C18 5 μm particles packed to 10 cm. Reverse-phase separation was performed on an Agilent 1100 using 0.1% acetic acid for solvent A and 0.1% acetic acid in 60% acetonitrile for solvent B for short peptides or 0.2% formic acid for solvent A and 0.2% formic acid in 80% acetonitrile for solvent B for longer peptides. Gradients were run from 0-100% solvent B in 60 minutes at a flow rate of 100 μL/min.

The preparatory columns were pressure loaded with 10 fmol equivalent of Pierce Retention Time Calibration Mixture and approximately 10-20 pmol of the digested material. The columns were rinsed with Solvent A prior to evaluation on a Thermo Orbitrap™ Fusion™ Mass Spectrometer. A full mass spectrum was acquired in the Orbitrap with 60,000 resolution at 200 m/z from 300 – 2000 m/z. Precursors were selected for fragmentation utilizing a decision tree process based on charge state and charge density (Figure S1F). Roughly, species with high charge density were fragmented by ETD; species with lower charge density were fragmented by CAD. Low-resolution MS/MS spectra were acquired for precursors with a charge state from +2 to +5, and high-resolution spectra were acquired for precursors with a charge state from +5 to +10. Precursors were excluded for 20 seconds after selection for fragmentation. Results were analyzed and manually validated using inhouse developed software for fragment masses.

### Preparative chaperone glutamate-glutamylation

Glutamylation of Npm2 and Nap1 proteins was performed as previously described ^9,30^. Briefly, 5 to 10mg Npm2 or Nap1 proteins (final concentration 2.5mg/mL) were mixed 1:100 with GST-TTLL4 (561-1199aa) in buffer containing 60mM Tris-HCl pH 8.5, 150mM NaCl, 10mM MgCl_2_, 2.5mM K+-L-Glutamate, 2.5mM ATP, and 5mM *β*-ME. Reactions were incubate at 30 °C for 2 h then another 1:100 aliquot of GST-TTLL4 was added and incubation continued for another 2 h; one additional spike-in of GST-TTLL4 was added and the reaction was incubated for another 4 h. Glutamylation was confirmed by immunoblot using the anti-Glu antibody.

### Competitive pull-down assays

For each sample, 10 μg H2A.S2/H2B dimers were mixed with 25 μL Streptactin Superflow resin in 200 μL binding buffer (50 mM Tris-HCl, pH 8.0, 150 mM NaCl, 5 mM *β*-ME) and incubated with rotation at 4 °C for 1 h. Resin only controls substituted binding buffer for H2A.S2/H2B. The resin was washed 3 × 500 μL using the same buffer. 2.5 μM chaperone proteins were added to the resin, either alone or mixed, in a final volume of 200 μL and incubated with rotation at 4 °C for 1h. Each sample was washed with binding buffer 8 × 200 μL; resin was transferred to a new tube after the 8^th^ wash and eluted with 25 μL 2x Laemmli Sample buffer. 10 μL of eluate was separated by 15% SDS-PAGE and gels were stained with Coomassie Brilliant Blue. All experiments were repeated at least twice; a representative experiment is shown.

### Histone capture disaggregation assay

H2A/H2B dimers were mixed at a 20:1 molar ratio with a 500 bp linear fragment of DNA (final concentration 400 nM histones and 20 nM DNA) in a 20 μL reaction volume in buffer containing 25mM sodium phosphate pH 7.0, 150mM NaCl, and 1mM EDTA. The reactions were incubated at 23 °C for 15 min. Npm2 or Nap1 proteins were added, and the reactions were incubated at 23 °C for an additional 30 min. 2.8 μL of 50% glycerol was added to each reaction, and 10 μL of each reaction was separated on 5% TBE native gel at 4 °C for 45 min at constant 150 V. The gels were post-stained with EtBr. All experiments were repeated at least twice; a representative experiment is shown.

### Tetrasome and mononucleosome assembly assays

Tetrasomes were assembled by salt dilution by mixing 187 bp DNA, containing the 147 bp Lowary and Widom 601 sequence ^5252^ flanked by 20 bp DNA on either end, with a 1.5-fold molar excess of (H3/H4)_2_ in a buffer containing 50 mM Tris-HCl pH 8.0, 2 M NaCl, 1 mM EDTA, 10 mM *β*-ME, and 10% glycerol. The mixture was incubated at 37 °C for 15 min then salt was diluted 10-fold (stepwise to 1.5, 1.0, 0.8, 0.7, 0.6, 0.5, 0.4, 0.3, and 0.2M NaCl) by the addition of salt-free buffer containing 50 mM Tris-HCl pH 8.0, 1 mM EDTA, 10 mM *β*-ME, 10% glycerol, 0.02% NP40, 1% PVA, and 1% PEG8K. The mixture was incubated at 37 °C for 15 min between each dilution.

Mononucleosome assembly assays were carried out by combining Npm2 or Nap1 proteins with H2A/H2B dimers at a 2-fold molar excess of the tetrasome concentration in a 25 μL reaction volume. Chaperone proteins and H2A/H2B were incubated at 30 °C for 15 min in the absence of tetrasomes. Tetrasomes were added and the reaction was incubated at 30 °C for an additional 15 min. 10 μL of the reactions were separated on 5% TBE native gels at 4 °C for 45 min at a constant 150 V, and gels were post-stained with EtBr. Nucleosomes assembled by salt dilution with HeLa octamers (as previously described ^3030^) were included on the gel as a sizing control. All experiments were repeated at least twice; a representative experiment is shown.

### Fluorescence Binding Assays

Intrinsic tryptophan fluorescence of the unmodified or glutamylated Npm2 A2 peptides (DYSWAEEEDE) in 20 mM Tris-HCl pH 7.6, 150 mM NaCl, 1 mM EDTA, 1 mM DTT, and 0.01% NP-40 was measured at 20°C in a Fluoromax-4 spectrofluorometer (HORIBA, Kyoto, Japan). The Npm2 peptide at 0.5 μM in a 0.1 × 0.1 cm quartz cuvette was excited at 295 nm using 5 × 5 nm slits. Emission spectra were collected in 1 nm steps between 305 nm and 450 nm using 0.5 s integration time per step. Background autofluorescence from buffer and histones was collected before addition of peptide and subtracted from spectra to obtain signal from the peptide alone. As previously described^99^, addition of histones resulted in a blue shift of the spectra, and data was plotted as the ratio of intensities at 340 nm and 360 nm (D340/360). Curves were converted to fractional saturation (Y) by normalizing D340/360_min_ and D340/360_max_ values to 0 and 1, respectively. Equilibrium dissociation constants (*K*_D_) and standard errors were then calculated by fitting the data to a single-site binding model taking the concentration of the peptide into account, as previously described ^99^:

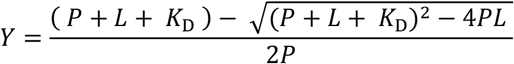

Where Y is the normalized response (0-1), P is the Npm2 peptide concentration (0.5 μM), L is the concentration of histone H2A/H2B dimers, and *K*_D_ is the equilibrium dissociation constant.

### Protein Thermal Shift Assays

All thermal denaturation assays used H2A/H2B dimers at 0.5 mg/mL in 50 mM Bis-Tris propane pH 7.4, 150 mM NaCl, 1 mM EDTA, 1 mM DTT, and 10X SYPRO Orange. Denaturation assays were performed in a 384-well plate in a qPCR machine (Roche Lightcycler II) using a temperature range of 25-99 °C, and fluorescence intensity was measured in 0.5 °C steps equilibrating for 10 seconds at each temperature prior to reading fluorescence. Reactions were excited at 495 nm and fluorescence emission was recorded at 600 nm. Fluorescence curves were blanked with buffer without H2A/H2B dimers. Chaperone peptides and DNA were added in slight excess at a 1.2:1 molar ratio. Fluorescence of chaperone peptides and DNA were also measured in the absence of H2A/H2B, and showed no difference compared to buffer alone. Each reaction was split and measured five times. Fluorescence intensities with errors from three replicates were fit to the equation:

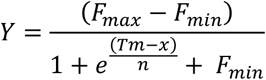

Where F_min_ is the starting fluorescence and F_max_ is the highest fluorescence intensity along the curve. T_m_ is the melting temperature, and *n* is the cooperativity coefficient. The three replicates (each composed of 5 technical replicates) were used to calculate final T_m_ values with standard error values, using the TSA-CRAFT algorithm^5353^. One-way ANOVA tests were performed to determine significance of ΔT_m_ values.

### Molecular dynamics simulations

All-atom, explicit solvent molecular dynamics (MD) trajectories were generated for the following four systems: (a) unmodified acidic IDR of Npm2 with H2A/H2B, (b) glutamylated acidic IDR of Npm2 with H2A/H2B, (c) unmodified acidic IDR of Nap1A3 with H2A/H2B, and (d) glutamylated acidic IDR of Nap1A3 with H2A/H2B. The Leap package implemented in AmberTools20 was utilized to parameterize the initial models of histone IDR/H2AH2B dimer^54^. The pdb4amber tool implemented in AmberTools20 was utilized to prepare the refined and optimized initial models to be used in subsequent Molecular Dynamics (MD) simulations. All MD simulations presented in this work were performed with Amber20 using Particle Mesh Ewald CUDA implementation (pmemd.cuda) and ff19SB force field^55^. All initial models were solvated in TIP4D water model with appropriate ions added to achieve neutrality. A solvent buffer of 15 Å was placed on all sides of each histone-peptide starting model. Periodic Boundary Conditions (PBCs) and Particle Mesh Ewald (PME) approximations for long-range interactions were implemented along with 9 Å cutoff for electrostatic interactions.

The implementation of minimization, equilibration, and production steps of all MD simulations consist of the following steps: (a) water and ions were minimized while other components were restrained with a positional restraint of 50.0 kcal/mol•Å^2^, (b) under constant volume, the system was slowly heated up from 100 K to 303.15 K over 10 ns of simulation time, (c) the system was kept at 303.15 K over 10 ns of simulation time at constant pressure, which allows the box density to relax, (d) keeping the system at 303.15 K at constant pressure, 5 ns of simulation was performed with a lower restraint of 10.0 kcal/mol•Å^2^, (e) minimization run was performed by applying restraint of 10.0 kcal/mol•Å^2^ on only the backbone atoms, (f) the system was relaxed over 10 ns of simulation time under constant pressure conditions with backbone atoms restrained with a coefficient of 10.0 kcal/mol•Å^2^, (g) the system was relaxed over 20 ns of simulation time under constant pressure conditions by lowering the restraint on the backbone atoms to 1.0 kcal/mol•Å^2^, and (h) the system was further relaxed over 20 ns of simulation time under constant pressure conditions by lowering the restraint on the backbone atoms to 0.1 kcal/mol•Å^2^. Following this extensive minimization and relaxation protocol, production simulations with no restraints were run in the NPT ensemble at 1.0 bar and 303.15 K with an integration timestep of 2.0 fs for a simulation length of 1 μs. The Langevin dynamics collision frequency was set to 1 ps^−1^ in production simulations. A unique random seed determined by the system clock was used for each Langevin dynamics simulation. Simulated trajectory frames containing solute and solvent atoms were collected at 10 ps intervals for subsequent analysis. Frames from the simulated trajectories were processed using the CPPTRAJ ^5656^ module in Amber20.

#### Determining intermolecular contacts between histone IDR and H2A/H2B dimer

Intermolecular atomic contacts between peptide IDR and histone H2A/H2B dimer were calculated from the simulated trajectory snapshots using INTERCAAT ^57^ using default settings.

#### Clustering of structural ensembles of simulated trajectory

To gain insight into the dominant conformations of IDPs a density-based conformational clustering approach was utilized ^58^, with a cutoff value of 1.25 Å.

#### Atomic fluctuation

Atomic fluctuation of histone dimer H2A/H2B (in the presence and absence of peptide IDR) was calculated by first computing an ensemble average structure. This average structure was structurally aligned using CPPTRAJ module of the AMBER20 package to the first reference frame, and RMSD for each residue was measured.

### Labeling Scheme and NMR Analysis of the Xenopus H2A/H2B Dimer

H2A and H2B were expressed individually, purified under denaturing conditions, and refolded into dimers as described previously ^49^. H2A and H2B were isotopically labeled using the Marley Method ^59^, except that M9 minimal media was made with 100% D_2_O for the ^13^C-^15^N-^2^H and ^15^N-^2^H labeled samples.

All NMR experiments were performed at the New York Structural Biology Center and Albert Einstein College of Medicine on Bruker Avance III 600 MHz, 700 MHz or 800 MHz spectrometers running TopSpin and equipped with 5 mm cryogenically cooled triple resonance probes or on a Varian INOVA 600 MHz spectrometer running VnmrJ and equipped with a 5 mm cryogenically cooled triple resonance probe. All NMR data sets were processed on NMRbox ^60^ with NMRPipe ^61^ and analyzed using CcpNmr Analysis Version 2 software ^62^. NMR samples were prepared in 25mM Na2PO4 pH 7.0, 150mM NaCl, 1mM EDTA, 10% D2O, 0.05mM TSP and run at 25° C. For peptide experiments, the final DMSO concentration was no more than 0.5% by volume. Control experiments on labeled histones showed no HSQC chemical shifts with 0.5% DMSO.

For assignment 0.6mM of either ^2^H-^13^C-^15^N H2A/H2B or H2A/^2^H-^13^C-^15^N H2B was used. Assignment of spectra from these two samples was accomplished by using a standard suite of triple resonance experiments (HNCO, HNcaCO, HNCA, HNcoCA, HNCACB, and HNcoCACB) utilizing TROSY-style pulse sequences and non-uniform sampling (NUS) methods ^63-65^ Resonances were assigned using a combination of PINE NMR server predictions and manual inspection of the spectra ^66^.

All CSP experiments were performed in an identical buffer by collecting TROSY-HSQC spectra of 0.3mM ^2^H-^15^N H2A/H2B or 0.3mM H2A/^2^H-^15^N H2B on a Bruker 600MHz spectrometer. Npm2 and Nap1 peptides were titrated by setting up multiple 50μL samples in 1.7 mm (about 0.07 in) NMR tubes and spectra were acquired using an autosampler. The Echo-Antiecho method with 2048 (t_2_) and 100 (t_1_) complex data points was used to acquire all the data, keeping the sweep width fixed at 14 and 32 ppm for ^1^H and ^15^N, respectively. A total of 64 scans and 32 dummy scans were recorded for each experiment.

Variable temperature experiments were performed by collecting TROSY-HSQC spectra using either 0.3mM ^2^H-^15^N H2A/H2B or 0.3mM H2A/^2^H-^15^N H2B on a single sample. The Echo-Antiecho method with 2048 (t_2_) and 128 (t_1_) complex data points was used to acquire all the data, keeping the sweep width fixed at 14 and 32 ppm for ^1^H and ^15^N, respectively. A total of 16 scans and 32 dummy scans were recorded for each experiment. After temperature change, the samples were equilibrated for 20 minutes at the new temperature prior to collecting a TROSY-HSQC spectrum. Peak intensity changes and CSPs were calculated as described previously ^9^.

DNA titration was performed using a 187 bp long DNA sequence comprising of a 147 bp core region from 601 Widom sequence flanked by 20bp at either end amplified from pGEM_3Z plasmid (ggtcgctgttcaatacatgcACAGGATGTATA-TATCTGACACGTGCCTGGAGACTAGGGAG-TAATCCCCTTGGCGGTTAAAACGCGGGGGACAGCGCGTACGTGCGTTTAAGCGGTGCTAGAGCTGTC-TACGACCAATTGAGCGGCCTCGGCACCGG-GATTCTCCAGggcggccgcgtatagggtcc). The DNA sample was also prepared by resuspending in NMR buffer as mentioned above. The Echo-Antiecho method with 2048 (T2) and 100 (T1) complex data points was used to acquire the data, keeping the sweep width fixed at 14 and 32 ppm for ^1^H and ^15^N, respectively. A total of 512 scans and 32 dummy scans were recorded for each experiment.

### Histone peptide arrays

Histone peptide array experiments were performed by JPT Peptide Technologies (Berlin, Germany) (Cat# His_MA_01). Binding events were probed by using a rabbit antibody raised against Nap1. Fluorescently labeled α rabbit IgG was used for detection of binding events. Signal intensities were scaled to the maximum intensity. Binding events on unmodified peptides were mapped onto a linear sequence of the histones.

## Supporting information

Supplementary Chemical Information

## Supporting Information

### Accession codes

Uniprot accesion numbers for proteins used in this study: *Xenoupus laevis* Nap1 (Q4U0Y4-2), Npm2 (A0A1L8GRP4), H2A (P06897), H2B (P02281), and human TTLL4 561-1199aa (Q14679).

Assigned chemical shifts for the *Xenopus laevis* H2A and H2B will be deposited in the Biological Magnetic Resonance Data Bank (BMRB) under accession (*pending*).

## Acknowledgments

This work was supported by the National Institutes of Health [R01GM135614 to D.S., R01GM037537 to D.F.H., R35GM136357 and R01AI141816 to A.F., T32GM007491 and F31GM116536 to C.W.]. D.S. was also supported for this project by the Irma T. Hirschl Trust. The Bruker 600 MHz NMR instrument in the Structural NMR Resource at the Albert Einstein College of Medicine was purchased using funds from NIH award 1S10OD016305 and is supported by a Cancer Center Support Grant (P30 CA013330). The data collected at NYSBC was made possible by a grant from ORIP/NIH facility improvement grant CO6RR015495. The 700 MHz spectrometer was purchased with funds from NIH grant S10OD018509. Data collected using the 800MHz Avance III spectrometer is supported by NIH grant S10OD016432. We are grateful to Anthony Spano and Anthony Frankfurter (Univ of Virginia) for the gift of the anti-glutamylation antibodies. We thank Michael Brenowitz for ongoing kibitzing and consultation and Sergei Khrapunov for assistance with fluorescence binding studies.

## Conflict of Interest

The authors declare that they have no conflicts of interest with the contents of this article.

## Author contributions

BML, CW, and HI conceived and executed biochemistry and NMR experiments, analysis, and wrote the paper. PN conceived, executed, and analyzed molecular dynamics studies; AF supervised molecular dynamics studies. SH performed biochemistry experiments and analyzed results. SC assisted with and analyzed NMR studies; DC supervised and analyzed NMR studies; SML, JS performed mass spectrometry analysis; JS and DFH supervised mass spectrometry studies. DS conceived and analyzed data, wrote the paper, and supervised this entire study. All authors read and approved the final manuscript.

To improve concision in portions of this manuscript, during the preparation of this work the author(s) used ChatGPT4.0. After using this tool, the authors fully reviewed and edited the content as needed and take full responsibility for the content of the publication.

## MD Simulation Movies

Supplementary Movies 1: Simulated molecular dynamics trajectories for histone dimer (H2A/H2B) with acidic IDR peptide for **a**. apo (no peptide); **b**. unmodified-Npm2 A2, **c**. glutamylated-Npm2 A2, **d**. unmodified-Nap1 A3, and **e**. glutamylated-Nap1 A3. The histones H2A and H2B are shown in gray, while the chaperone acidic IDR peptides, Npm2 A2 and Nap1 A3 are shown in orange. Glutamates and glutamyl-glutamates are shown with sidechains with transparent surfaces. The starting conformation between the histone dimer and peptide chaperones was spatially separated by at least 20 Å.

## Supplementary chemical information

PDB file representing **a**. Glutamate (glu) and **b**. γ-glutamyl-glutamate (Eglu)

Schrödinger Maestro project files for **c**. Glutamate (glu) and **d**. γ-glutamyl-glutamate (Eglu)

***Extended Discussion (pages S2-S3)***

***Supplementary Figures S1-S6 (pages S4 – S9)***

***Supplementary References (page S10)***

***Supplementary Table S1-*** This spreadsheet contains the NMR intensity measurements from ^2^H-^15^N-H2A/H2B heterodimers interacting with the Npm2 Nap1 IDR peptides.

***Supplementary Table S2-*** This spreadsheet contains the NMR chemical shift perturbations from ^2^H-^15^N-H2A/H2B heterodimers interacting with the Npm2 Nap1 IDR peptides.

***Supplementary Table S3-*** This spreadsheet contains the intensity measurements from H2A/^2^H-^15^N-H2B heterodimers interacting with the Npm2 Nap1 IDR peptides.

***Supplementary Table S4-*** This spreadsheet contains the NMR chemical shift perturbations from H2A/^2^H-^15^N-H2B heterodimers interacting with the Npm2 Nap1 IDR peptides.

## EXTENDED DISCUSSION

The structural models from the previous NMR study of the human H2A/H2B dimer in solution show a high positional RMSD between calculated conformers ^1^. More specifically, they observed a large amount of flexibility at the H2A α1 helix (residues 28-41) and the H2B C-terminal helix (residues 104-125). Even within the remainder of the histone fold positional RMSDs of their top ten conformers ranged from ∼1-5Å, and measurements such as NMR HDX, HET^ex^-BEST-TROSY, and heteronuclear NOE experiments suggest that the H2A/H2B dimer is highly dynamic in solution. Our NMR analysis of the *Xenopus* H2A/H2B dimer supports this idea of a dynamic H2A/H2B dimer in solution. Particularly, the global loss of peak intensities for all residues in the histone fold upon both temperature reduction and chaperone peptide binding likely indicates that this is an effect of the stabilization of the H2A/H2B dimer.

We suspect that the global loss of peak intensities in the histone fold upon glutamylated chaperone peptide binding reflects a dramatic change in conformational dynamics upon chaperone binding. Residues from the histone fold may be undergoing fast conformational exchange when free in solution, and that chaperone binding may induce peak disappearance by stabilizing the histone fold, thus slowing this exchange process into the intermediate/slow regime. In support of this, previous hydrogen-deuterium exchange (HDX) experiments of the H2A/H2B dimer showed that most amide protons in the histone fold exchange rapidly with a deuterium-based buffer, indicating it is highly dynamic in solution ^1,2^.

We also observed that the αC helix of H2A is slightly more extended in the unbound state compared to nucleosomal H2A/H2B. Extension of this helix would likely form clashes with other histones in the nucleosome. It’s also worth noting the H2A.Z/H2B chaperones ANP32E and YL1, which both specifically remove H2A.Z/H2B dimers from nucleosomes. Structures of acidic peptides from these chaperones bound to H2A.Z/H2B dimers show a similar extension of the αC of H2A.Z ^3-5^. The authors of these studies suggest that this helix extension is critical for the removal of H2A.Z/H2B dimers from the nucleosome, and that its presence would prevent the chaperone bound dimers from being incorporated. Taken together, the loss of the αN helix and the extension of the αC helix in solution suggest that major conformational changes must occur in H2A for H2A/H2B dimers to be incorporated into nucleosomes. However, given that we observe similar αC extension in solution, it remains unclear whether ANP32E and YL1 binding induce the extension of this helix or if they preferentially bind to the extended form. Overall, this may suggest a generalizable mechanism for chaperones, which may either induce these conformational changes thus promoting deposition of H2A/H2B or prevent these conformational changes from occurring thus promoting the removal of H2A/H2B from nucleosomes.

An interesting observation in this study was the reappearance of certain residues peak like S55 and S60 in the Nap1 A3glu spectra, a phenomenon referred to as peak rescue, which had completely broadened upon addition of non-glutamylated Nap1 A3 peptide. This peculiar behavior was more profound for peptide interaction with H2A/ndH2B dimer. Complete line broadening upon ligand addition happens due to fast/intermediate chemical exchange processes and other factors that alter the local chemical environment of the nuclei in consideration. Reappearance of such peaks upon titration with Nap1A3glu peptide could be an indication of multiple conformational states or binding modes depending on the ligand. In any case, such rescue phenomenon indicates that the glutamylated peptide can make better contacts with the histone dimers due to higher affinity.

Our MD and NMR data show that IDR glutamylation enhances these localized interactions and leads to long-range stabilization of the histone fold. There are a few possible stabilization mechanisms: 1) localized acquisition of helices, such as the αN and 3_10_ on either end of H2A that are known to be present along the DNA binding interface in the nucleosome, to promote specific DNA interactions; 2) augmented electrostatic interactions between the glutamylated acidic IDRs and histones might set up a network of localized interactions that propagate across the histone molecules, culminating in a stabilized histone fold. This stabilization serves a dual role: 1) It facilitates the capture and orientation of free histones, thus aiding in nucleosome assembly, and 2) it assists in capturing histone aggregates from DNA, thereby acting as a ‘molecular rake’ to clear histone:DNA aggregates to accommodate proper nucleosome formation. In this way, glutamylation of Npm2 and Nap1 acidic IDRs improves both nucleosome assembly and the stability of newly deposited histones.

Histone dimer H2A/H2B and apo and glutamylated peptides are separated by at least 20 Angstrom. In our simulations, we observe that both apo and glutamylated peptide migrate to the histone surface within a timescale of ∼ 50 ns (during the initial phase of the MD trajectory), presumably guided by interactions between +*ve* charged flanking LYS residues on H2A/H2B and -*ve* charged glutamate on peptide IDR. Once the initial “encounter complex” is established, the glutamylated peptide forms a much more stable ensemble compared to the apo peptide. This demonstrates that the glutamylated peptide forms a “specific” binding pocket that is stable during the extended MD run, while the apo peptide samples a broader range of residues resulting in a binding pocket being transiently “non-specific”. It is essential that we extend the simulation up to a significant timescale because that buttresses the overall stability of the histone dimer:IDR bound ensemble in the glutamylated state compared to the apo state. The formation and subsequent stability of the initial encounter complex following equilibration emphasizes the observation that the coordination of chaperone IDR is significantly higher for glutamylated peptide compared to the apo peptide.

Considering other modifications on the histone chaperones suggests additional regulatory possibilities. Our previously identified arginine methylation on Npm2 C-terminal basic patch ^6^ may play a role in RNA-binding protein storage and release during early embryogenesis ^7^. Here, we identified farnesylation on the Nap1 X1 variant. Future studies may uncover possible mechanisms by which this PTM anchors the chaperone on the nuclear or plasma membranes. One hypothesis is that there is a cell cycle or developmentally regulated release of Nap1, coupling histone storage and release to key biological transitions.

**Supplementary Figure S1.**
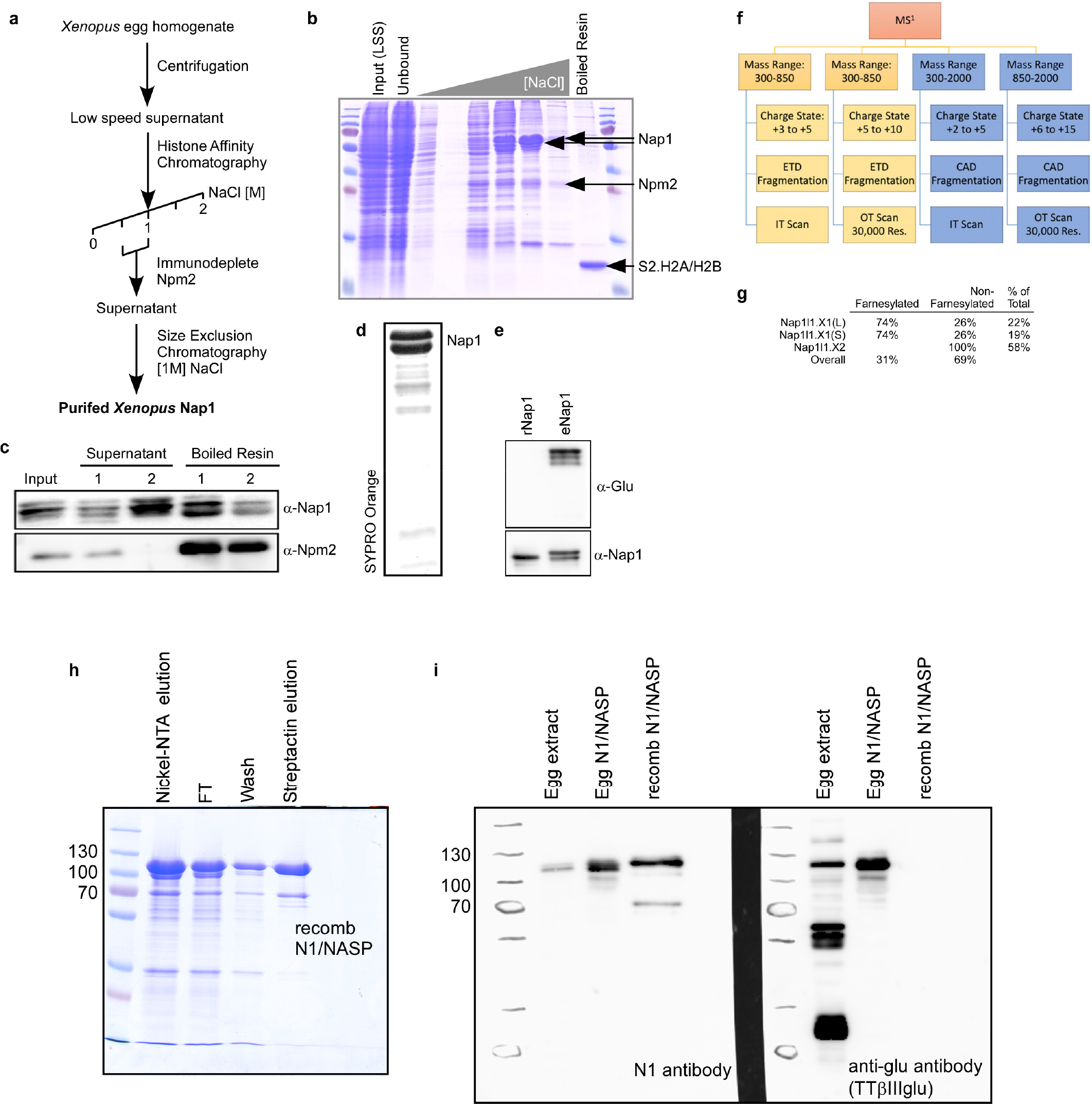
(Related to Figure 1). Characterization of post-translational glutamylation on *Xenopus laevis* histone chaperones. **a**. Chromatography scheme for purification of *Xenopus laevis* egg Nap1. **b**. Coomassie stained gel showing purification of Nap1 **c**. Immunoblot of Npm2 clearance step showing specific flow-through of Nap1 relative to Npm2, which remained on the resin **d**. SYPRO Orange protein stain of final Nap1 purified product. Two bands represent the predominant genetic isoforms, Nap1.L and Nap1.S **e**. Glutamylation immunoblot showing that recombinant (rNap1) is not glutamylated but purified egg (eNap1) is post-translationally glutamylated. Bottom blot shows general Nap1 antibody staining. **f**. Decision tree approach for mass spectrometry analysis of egg Nap1 **g**. Table showing identified Nap1 isoforms and the percentage of each isoform that are post-translationally farnesylated **h**. Purification of recombinant *Xenopus laevis* NASP (originally known as N1) **i**. Immunoblot showing egg extract, purified egg NASP, and recombinant NASP, blotted on left with a NAS/N1 antibody, and on the right with anti-glu antibody, demonstrating that NASP is also post-translationally glutamylated.

**Supplementary Figure S2.**
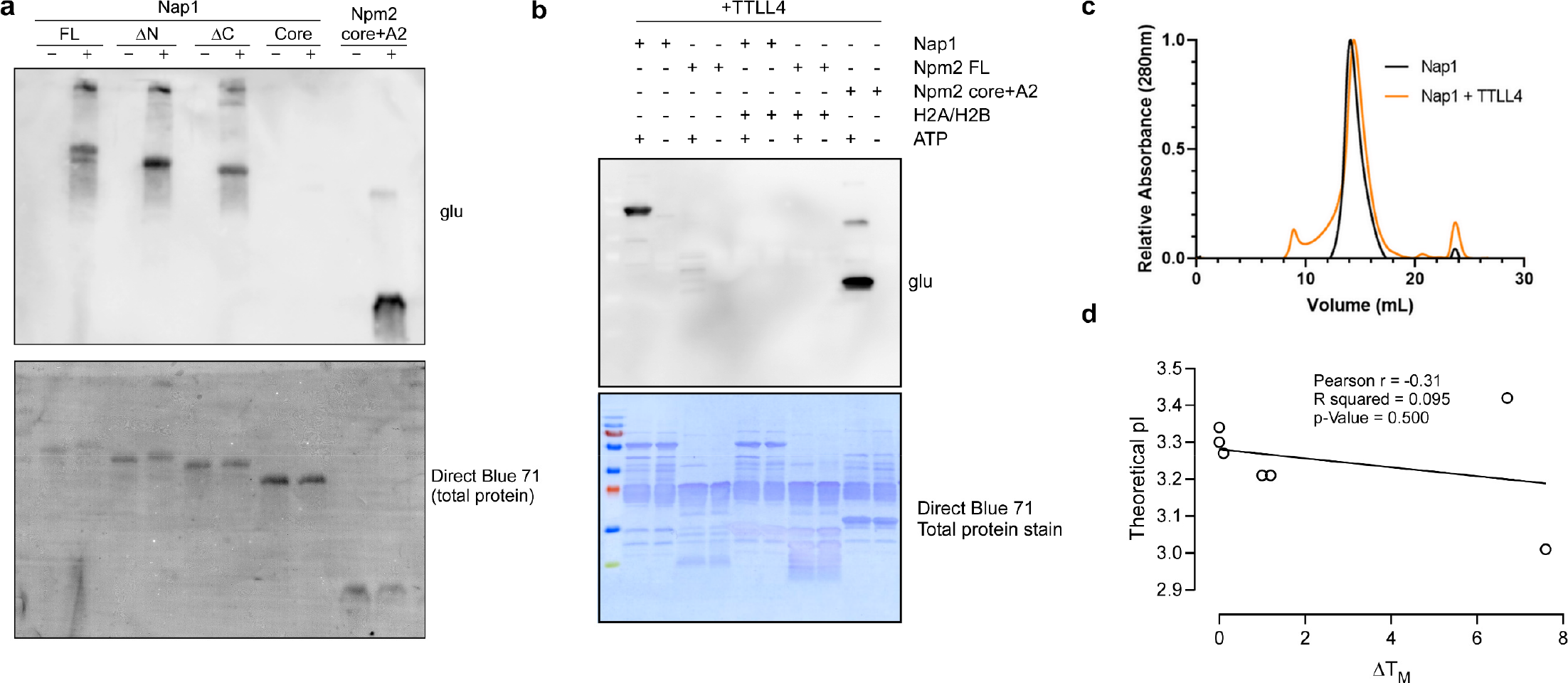
(Related to Figure 3). Glutamylation characterization. **a**. TTLL4 treatment of Nap1 N- and C-terminal truncations revealed that both N- and C-terminal acidic IDRs are post-translationally glutamylated. Top: anti-glu immunoblot; Bottom: Direct Blue 71 total protein stain. Npm2 core+A2 is shown as a positive control **b**. Histone chaperones Nap1, Npm2 full length (FL), and Npm2 core+A2 incubated with TTLL4 enzyme and with or without H2A/H2B or with or without ATP (to promote TTLL4 catalysis) were immunoblotted for glutamylation (glu) top and stained for total protein (bottom). The presence of histones blocked glutamylation of both Nap1 and Npm2. **c**. Nap1 oligomerization state was not impacted by post-translational glutamylation as compared to Nap1 alone (black), TTLL4 treated Nap1 (orange) did not change its elution profile on a Superose 6 gel filtration column. is also post-translationally glutamylated. **d**. Correlation plot between the calculated theoretical isoelectric point (pI) and change in melting temperature (ΔT_m_) of tested polyelectrolytes.

**Supplementary Figure S3.**
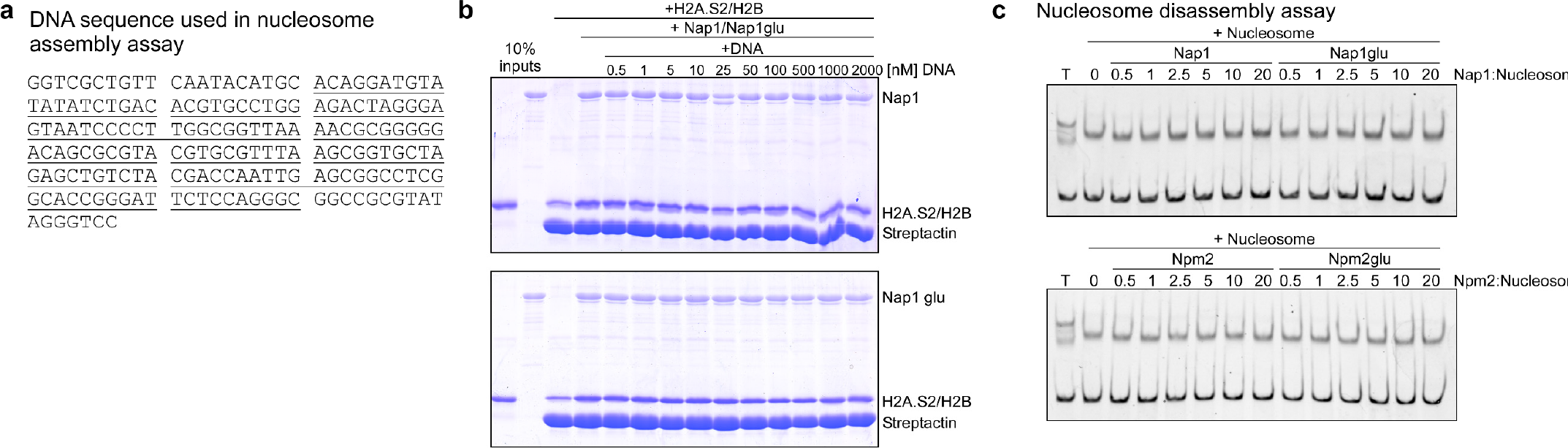
(Related to Figure 4. Chaperone glutamylation consequences. **a**. DNA sequence used in mononucleosome assembly assay. Underlined sequence = Widom 601’ **b**. StrepII-tagged H2A.S2/H2B pulldown binding assays of Nap1 (top gel) and TTLL4 treated Nap1 (Nap1glu, bottom gel) were competed with increasing concentrations of dsDNA (molarity as indicated). Coomassie stained gel with components included as indicated at the top. DNA did not outcompete chaperones from the histones. **c**. Mononucleosomes assembled by salt dialysis were treated with increasing amounts of Nap1 and Nap1glu (top gel) or Npm2core+A2 or Npm2core+A2glu (bottom gel) and visualized on a polyacrylamide gel stained with ethidium bromide. Chaperones did not disassemble mononucleosomes.

**Supplementary Figure S4.**
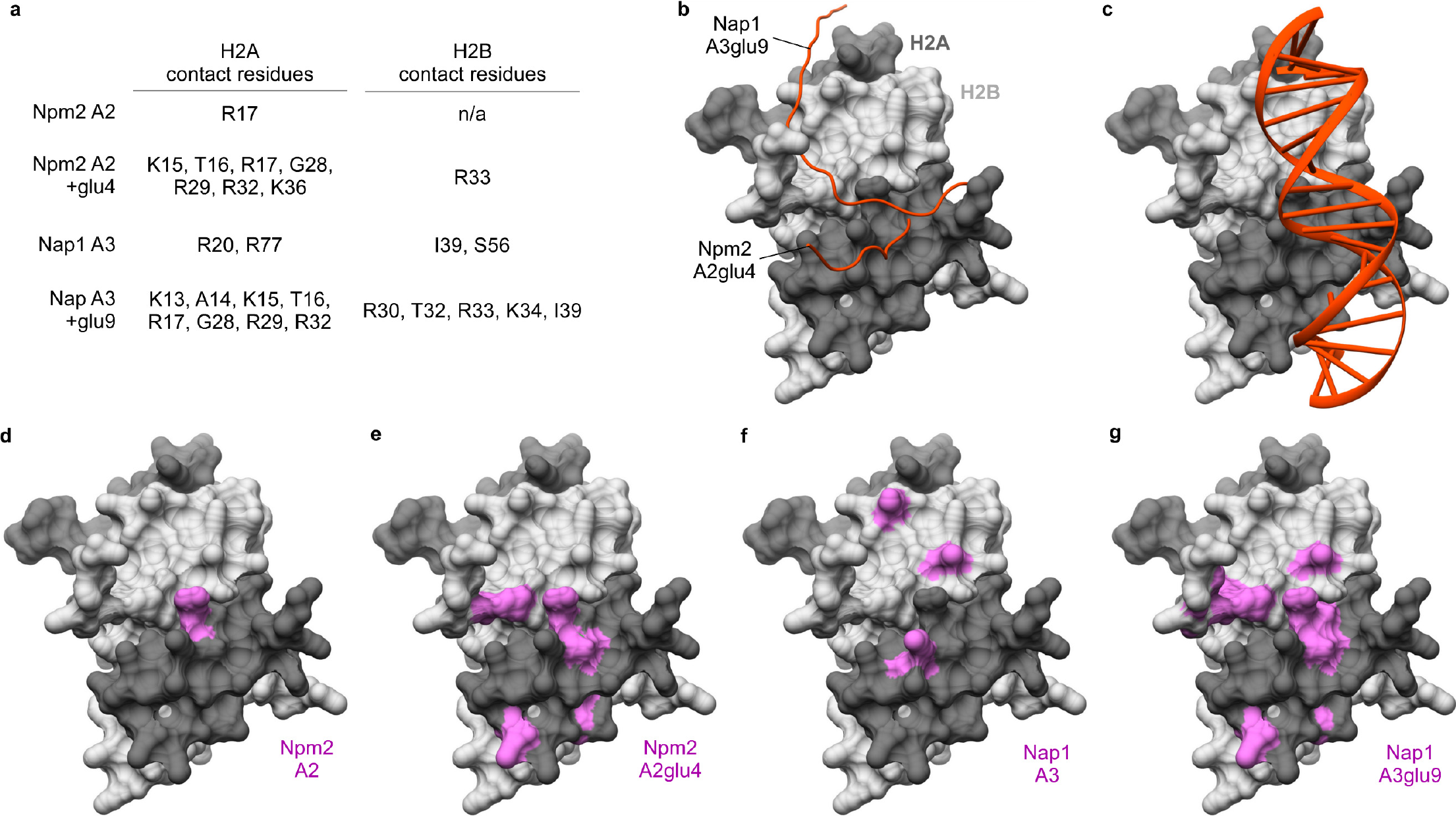
(Related to Figure 5). Models of molecular dynamic simulations. **a**. H2A and H2B residues in contact with apo and glutamylated peptide for the dominant clustered conformation. **b**. Dominant conformations for glutamylated Npm2A2 and Nap1A3 peptide. **c**. DNA (red) from nucleosome core particle (PDB:1AOI) shown on the surface of H2A/H2B (dark gray/light gray), chains C,D **d**. Positions in histones H2A/H2B (pink) that associate (<3.5Å average distance) with Npm2A2 during simulation **e**. Positions in histones H2A/H2B (pink) that associate (<3.5Å average distance) with Npm2A2glu4 during simulation **f**. Positions in histones H2A/H2B (pink) that associate (<3.5Å average distance) with Nap1A3 during simulation **g**. Positions in histones H2A/H2B (pink) that associate (<3.5Å average distance) with Nap1A3glu9 during simulation

**Supplementary Figure S5.**
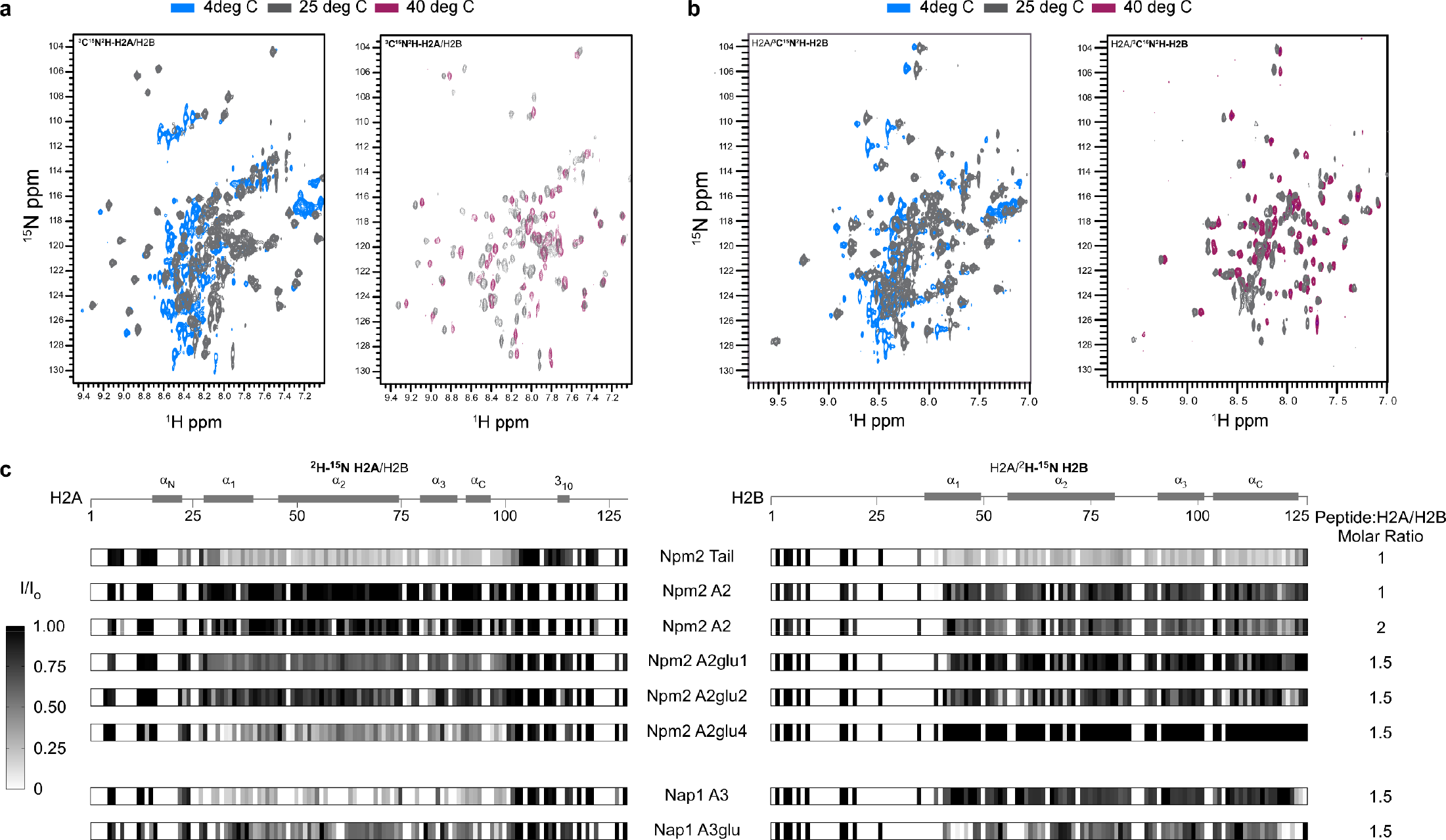
(Related to Figure 6). NMR analysis of histone chaperone binding to H2A/H2B. **a**. Two-dimensional overlay of the ndH2A/H2B HSQC spectra at 4°, 25°, and 40° C, respectively, displaying the change in peak intensities and spectral characteristic for nearly all residues within the histone core. VT-NMR experiments were recorded on Bruker Avance III 600 MHz using 0.3mM ^2^H-^15^N H2A/H2B samples. **b**. Two-dimensional overlay of the H2A/ndH2B HSQC spectra at 4°, 25°, and 40° C, respectively, displaying the change in peak intensities and spectral characteristic for nearly all residues within the histone core. VT-NMR experiments were recorded on Bruker Avance III 600 MHz using 0.3mM H2A/^2^H-^15^N H2B samples. **c**. Heat map depicting the consequence of peptide glutamylation upon histone binding, measured by plotting the global changes in peak intensity (I/I_0_) after titration of indicated peptides into ^2^H-^15^N H2A/H2B (left), and H2A/^2^H-^15^N H2B (right) sample. Compared to their non-glutamylated variants, the glutamylated peptides (both Npm2 A2 and Nap1 A3) depicted a much larger line broadening effect indicating stronger binding affinity. Additionally, Nap1 A3glu displayed more profound changes than Npm2 A4glu. The experiments were recorded on Bruker Avance III 600 MHz using 0.3mM ^2^H-^15^N H2A/H2B or H2A/^2^H-^15^N H2B samples.

**Supplementary Figure S6.**
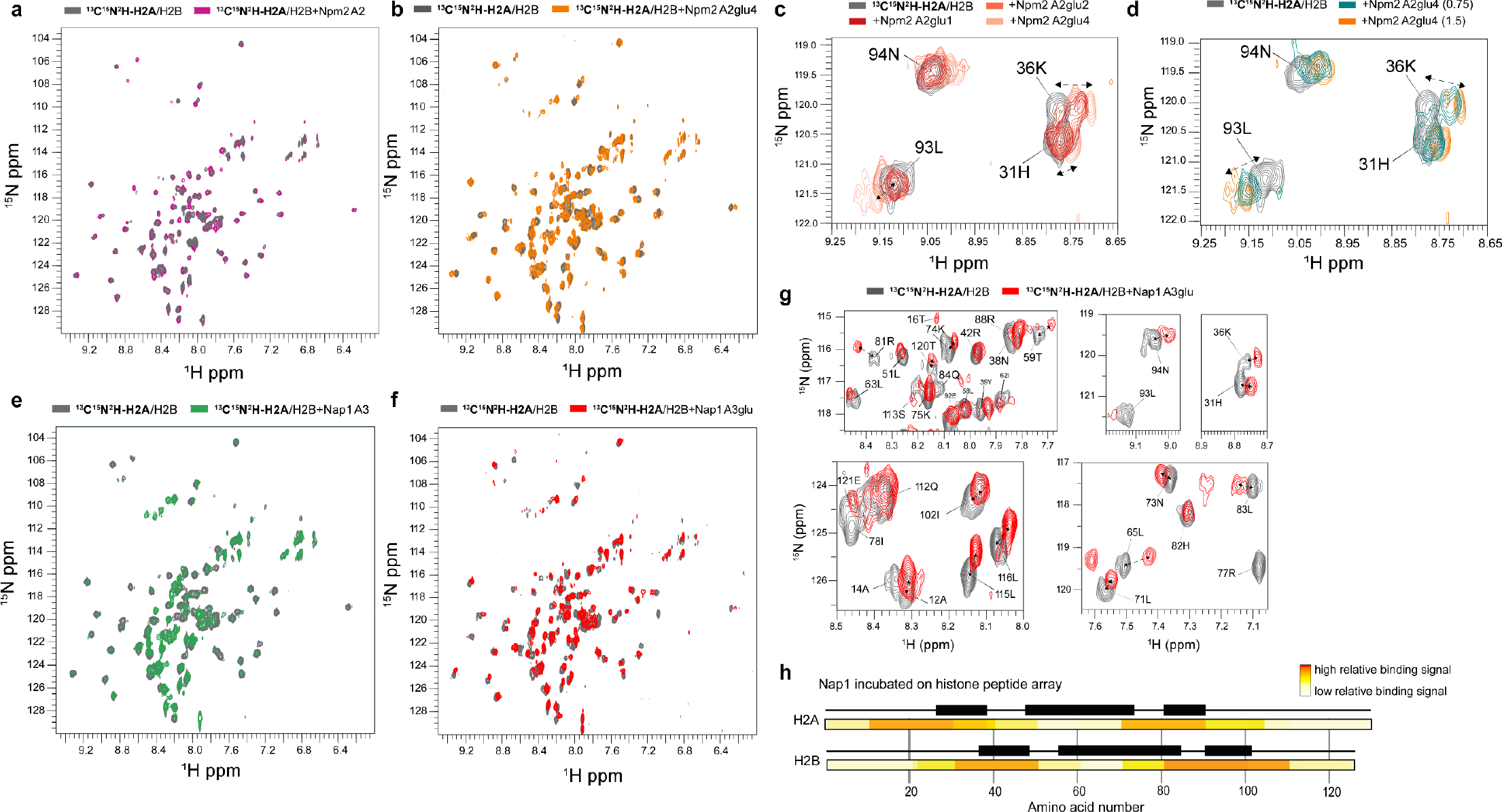
(Related to Figure 7). NMR analysis of histone chaperone binding to H2A/H2B. **a**. Two-dimensional overlay of the ^2^H-^15^N-H2A/H2B HSQC spectra upon titration with Npm2 A2 peptide. The experiments were recorded on Bruker Avance III 600 MHz using 0.3mM ^2^H-^15^N H2A/H2B at 25° C **b**. Two-dimensional overlay of the ^2^H-^15^N-H2A/H2B HSQC spectra upon titration with Npm2 A2glu4 peptide. The experiments were recorded on Bruker Avance III 600 MHz using 0.3mM ^2^H-^15^N H2A/H2B at 25° C **c**. Two-dimensional overlay of the ^2^H-^15^N-H2A/H2B HSQC spectra depicting the importance of glutamate-glutamylation for Npm2 A2 peptide interaction with histones. At similar, histone dimer to peptide ratio (1:1.5), Npm2 A2glu4-DYSWAE^E^E^E^E^E^DE^E^, with 4 consecutive glutamylation depicted significantly higher levels of spectral perturbations. The experiments were recorded on Bruker Avance III 600 MHz using 0.3mM ^2^H-^15^N-H2A/H2B at 25° C. **d**. Two-dimensional overlay of the ^2^H-^15^N-H2A/H2B HSQC spectra depicting the concentration-dependent increase in line broadening and/or CSP effects upon titrating with the glutamylated synthetic peptides Npm2. Shown are some of the key residues predicted to be involved in the interaction with histones. The experiments were recorded on Bruker Avance III 600 MHz using 0.3mM ^2^H-^15^N H2A/H2B at 25° C. **e**. Two-dimensional overlay of the ^2^H-^15^N-H2A/H2B HSQC spectra upon titration with Nap1 A3 peptide. The experiments were recorded on Bruker Avance III 600 MHz using 0.3mM ^2^H-^15^N H2A/H2B at 25° C. **f**. Two-dimensional overlay of the ^2^H-^15^N-H2A/H2B HSQC spectra upon titration with Nap1 A3glu peptide. The experiments were recorded on Bruker Avance III 600 MHz using 0.3mM ^2^H-^15^N H2A/H2B at 25° C. **g**. Two-dimensional overlay of the ^2^H-^15^N-H2A/H2B HSQC spectra depicting line broadening and/or CSP effects upon titrating with the glutamylated synthetic peptides Nap1 A3glu at 1:1.5 histone dimer to peptide ratio. Shown are some of the key residues predicted to be involved in the interaction with histones. The experiment was recorded on Bruker Avance III 600 MHz using 0.3mM ^2^H-^15^N H2A/H2B at 25° **h**. Peptide array studies with H2A/H2B dimers depicted higher binding affinity for residues in similar regions as predicted by NMR and MD simulation analysis, both for full length Npm2 and Nap1.

## Notes

### Competing Interest Statement

The authors have declared no competing interest.

